# The conserved bridging domain on HCV E1E2 glycoprotein complex is targeted by neutralizing antibodies from diverse lineages

**DOI:** 10.1101/2025.11.05.686883

**Authors:** Fang Chen, Yen Thi Kim Nguyen, Yi-Zong Lee, Erick Giang, Sing Chau Lau, Yukiye Koide, Shr-Hau Hung, Lynn Ueno, Linling He, Thomas R. Fuerst, Georg M. Lauer, Robyn L. Stanfield, Jiang Zhu, Ian A. Wilson, Mansun Law

## Abstract

The induction of potent and cross-reactive neutralizing antibody (nAb) responses remains a challenge in vaccine development against antigenically diverse viruses such as hepatitis C virus (HCV). The HCV E1E2 glycoprotein complex contains two major neutralizing sites: the neutralizing face (NF) and the less explored bridging domain (BD). Here, we characterized 25 BD-targeting nAbs isolated from infection or immunization. These antibodies arise from diverse B cell lineages but share convergent CDRH3 features. Epitope mapping by alanine scanning and negative-stain electron microscopy revealed overlapping epitopes on BD spanning antigenic regions AR4 and AR5, with variable back layer engagement. The crystal structure of a non-human primate BD nAb RM3-26 in complex with E2 uncovered a back layer-directed recognition mode analogous to that of the human nAb hcab40. Together, BD- and NF-directed nAbs exhibited additivity in their neutralization, highlighting BD as a conserved site of vulnerability on HCV and a valuable target for rational vaccine design.

## Introduction

Elicitation of neutralizing antibodies (nAbs) capable of cross-neutralizing a broad spectrum of viral isolates, i.e., broadly nAbs (bnAbs), is a central goal of rational vaccine design against antigenically diverse pathogens such as hepatitis C virus (HCV). Thus far, clinical trials of HCV vaccine candidates have failed to elicit robust immune responses with sufficient breadth to protect across diverse genotypes.^1,2^ This failure reflects multiple challenges, including viral genetic variability, glycan shielding of neutralizing epitopes, and inefficient activation of multifunctional T cells and bnAb precursor B cells by current immunogens.^3–5^ Although cellular immunity has long been regarded as the principal determinant of HCV control, accumulating evidence demonstrates that humoral immunity, particularly bnAb responses, also confers protection against HCV infection in humans and surrogate animal models.^6,7^ The discovery of bnAbs underscores the capacity of the immune system to find solutions to defend against antigenically diverse viruses.^1,8–11^ A deep understanding of bnAbs and their conserved epitopes will be critical for guiding the rational design of effective HCV vaccines.

HCV envelope glycoproteins E1 and E2 are natural targets for HCV bnAbs. Together, they form a non-covalent heterodimer on the viral surface.^12,13^ E2 interacts with key receptors on host cells, including tetraspanin CD81 and scavenger receptor class B type I (SR-BI), to initiate viral entry, while E1 is implicated in membrane fusion.^14,15^ Epitope mapping and structural characterization of numerous HCV bnAbs have identified two major, conserved sites of vulnerability on E1E2. The first is the E2 neutralizing face (NF), a hydrophobic surface comprising the E2 front layer and the CD81 binding loop (CD81bl) regions.^16^ This surface includes a major discontinuous antigenic region AR3, as well as two continuous antigenic sites, AS412 and AS434. AR3 is the dominant target of bnAb responses elicited during both natural infection and immunization.^1,11,17,18^ AR3-specific bnAbs, such as AR3A/B/C/D and HEPC3/74, demonstrate a strong bias toward derivation from the human immunoglobulin heavy chain variable gene fragment *IGHV1-69*.^19^ The second conserved site of vulnerability is the bridging domain (BD) on the E1E2 complex. The BD spans E2 residues 646-704 but its native conformation requires the presence of E1.^12,20^ BD contains two known non-overlapping antigenic regions, AR4 and AR5.^21^ Cross-nAb responses specific to BD are considered subdominant as most bnAbs elicited by wild-type E1E2 immunization^1^ or natural infection^22^ primarily target the E2 NF. Recognition of the BD requires an intact E1E2 heterodimer, underscoring the quaternary structural dependence of these epitopes.^12,20^ To date, only a small number of BD-targeting nAbs (BD nAbs), i.e., AR4- or AR5-binding antibodies, have been described in HCV-infected individuals (e.g., AR4A, AR5A, AT1618, HEPC111 and HEPC130),^21,23,24^ or in E1E2-immunized non-human primates (NHPs) (e.g., RM3-26 and RM7-22).^1^ Their apparent scarcity may stem from the technical challenges of studying the natively folded E1E2 complex.

Recent advances in the development of soluble, secreted recombinant E1E2 proteins have greatly facilitated the isolation and characterization of BD-targeting antibodies.^20,25^ Using these proteins, we recently identified a new panel of nAbs from people living with HCV (PLWHCV),^22^ of which 20 were BD-directed. Here, we set out to characterize the molecular and functional properties of this important antibody class by integrating genetic, functional, and structural analyses of these newly isolated and previously discovered BD nAbs. We show that BD nAbs display diverse lineages but often share recurring heavy chain complementarity determining region 3 (CDRH3) motifs, indicative of convergent evolution in epitope recognition. Alanine scanning mutagenesis and negative-stain electron microscopy (nsEM) revealed overlapping epitopes on BD across AR4 and AR5, with varying engagement of the E2 back layer region. RM3-26, a representative BD nAb, binds an epitope spanning the BD and back layer with weak affinity for soluble E2. The crystal structure of RM3-26-E2 core complex defined the molecular basis of this interaction. Furthermore, combinations of BD- and NF-targeting nAbs exhibited enhanced neutralization breadth and potency, highlighting the benefit of vaccine strategies that co-targeting these complementary antigenic sites. Together, these findings delineate the molecular and immunogenetic features of BD-targeting bnAbs, providing a structural framework to guide rational HCV vaccine design.

## Results

### Identification of HCV nAbs targeting conserved epitopes on E1E2 BD

We recently isolated a panel of HCV monoclonal antibodies (mAbs) from PLWHCV using a soluble E1E2 antigen probe.^22^ A subset of the mAbs (n = 20; Table S1) preferentially recognized HCV-1 SpyE1E2 (a soluble E1E2 comprising E1 residues 192-354 and E2 residues 384-717 fused to a SpyTag/SpyCatcher scaffold;^26^ Table S2), while showing low affinity or no detectable binding to soluble E2 ectodomain (E2ecto, residues 384-645;^27^ Table S2), which lacks the BD region (Figures 1A, left two panels, and S1A). We conducted detailed genetic, structural, and functional analyses of these antibodies together with five previously reported antibodies with similar binding properties, including AR4A,^12,20,21^ AR4B,^21^ AR5A,^21^ RM3-26,^1^ and RM7-22.^1^ Among 25 antibodies analyzed, 16 (64%) neutralized all six selected viral strains with tier 1 or tier 2 antibody resistance in the HCV pseudoparticle (HCVpp) panel, and 10 of these (40%) also neutralized at least three of seven tier 3-4 resistant strains (Figure 1A, third panel).

**Figure 1.**
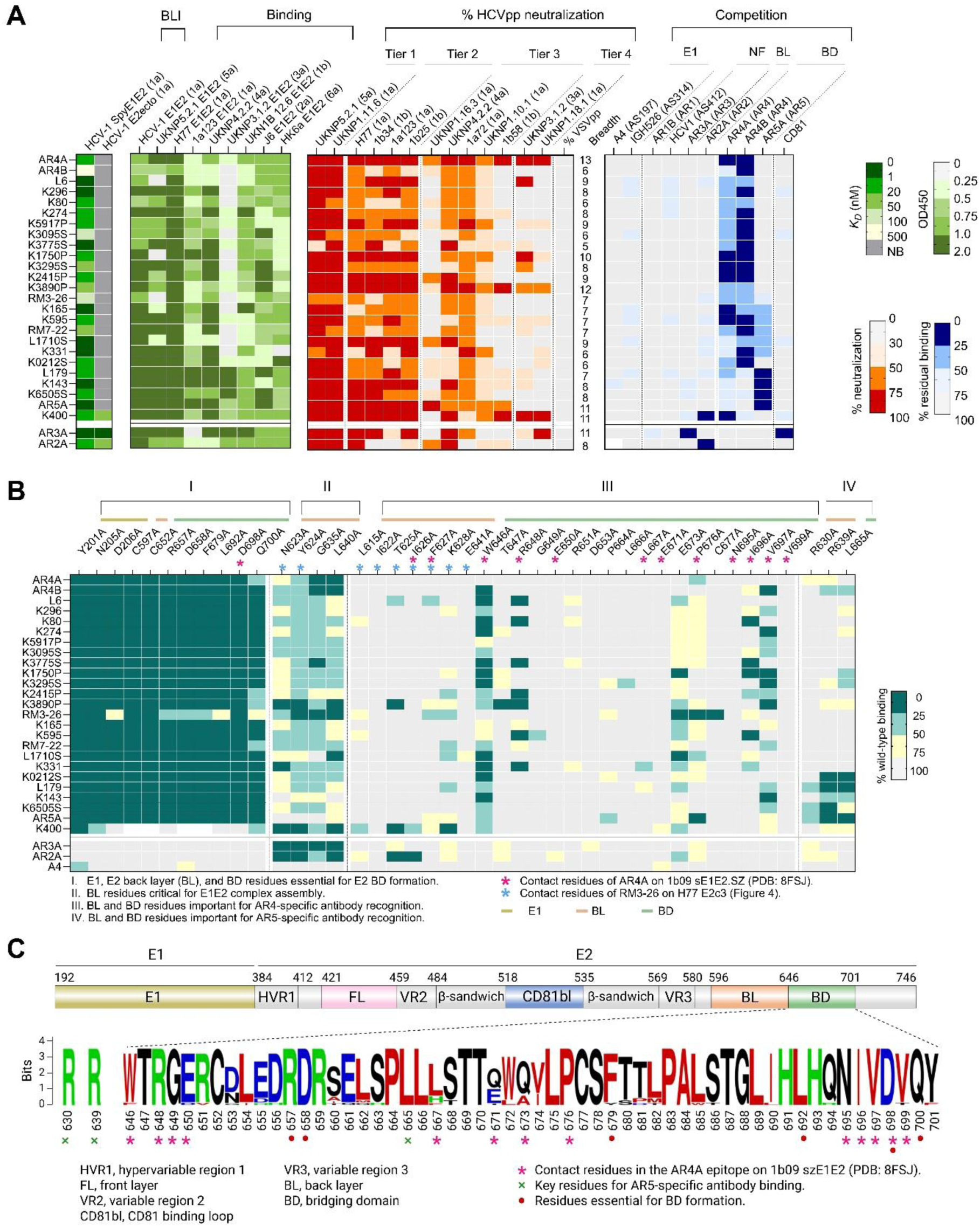
Function and specificity of HCV BD nAbs. (A) Binding affinity, cross-binding, neutralizing activity, and epitope specificity of BD-targeting bnAbs, assessed by biolayer interferometry (BLI), binding ELISA, HCVpp neutralization, and competition ELISA, respectively. Antibodies were tested at single concentrations of 50 µg/mL in BLI, 1 µg/mL in the binding ELISA, 25 µg/mL in HCVpp neutralization, and 20 µg/mL the competition ELISA. NF, neutralizing face. BL, back layer. Human nAbs AR3A (targeting the E2 NF) and AR2A (targeting the E2 back layer) were included as controls. (B) Epitope mapping of BD nAbs by alanine-scanning mutagenesis. Single alanine substitutions were introduced at selected residues of H77 E1E2. Antibody binding to mutant proteins was assessed by ELISA, and results are presented as a heatmap showing binding relative to wild-type (WT) H77 E1E2. Human nAbs AR3A and AR2A, as well as mouse antibody A4 were included as controls. (C) Schematic of E1E2 subregions and sequence logo of the BD region showing amino acid conservation across 18 HCV strains used in ELISA and neutralization assays (Figure S2).

Competition ELISA with reference human mAbs or CD81 revealed three major binding patterns (Figure 1A, right panel). The majority of antibodies, including L6, K274, K296, and K05917P, competed with antibodies AR4A and AR4B, implicating AR4 as their dominant target. A subset (L179, K143 and K6505S) competed with AR5A, defining AR5 specificity. Several antibodies, such as K165 and K595, competed with both AR4A and AR5A, indicating recognition of a composite AR4/AR5 surface. Notably, antibody K400 competed with both AR4A and AR2A (a human antibody recognizing a conformational epitope on the E2 back layer independent of E1), suggesting its epitope spanning BD and back layer. None of these antibodies competed with CD81, indicating that BD epitopes are located outside the receptor-binding site (Figure 1A, right panel). To dissect the contribution of individual residues on E1E2 to antibody recognition, we examined the binding of BD antibodies to E1E2 alanine mutants at selected positions in E1, back layer, and BD (Figure 1B). Mutations at Y201, N205, and D206 (E1); C597 (E2 back layer); and C652, R657, D658, F679, L692, D698, and Q700 (E2 BD) completely abrogated binding of nearly all BD antibodies but did not affect antibodies targeting other epitopes, such as AR3A,^17,28^ AR2A(recognizing a discontinuous epitope on the E2 back layer),^17^ or A4 (recognizing an linear epitope on E1)^29^ (Figure 1B). These results highlight the essential role of these residues in maintaining BD conformation. In contrast, mutations at N623, Y625, G635, and L640 in the E2 back layer reduced binding of all antibodies recognizing conformational epitopes (e.g., BD antibodies, AR3A and AR2A) but not the linear epitope-targeting antibody A4, suggesting a role in global E2 folding and structural integrity rather than direct epitope contact. These findings are consistent with previous reports.^30^ Further analysis of AR4 and AR5 epitopes showed that BD residues W646, R648, E673, P676, I696 and V697 were essential for binding by most AR4-specific antibodies, whereas R630, R639 (back layer), and L665 (BD) were indispensable for AR5-specific antibodies (Figure 1B). Moreover, back layer residues such as T625, F627 and K628 were required for recognition by certain BD nAbs, such as K3890P, RM3-26, and K400, reflecting antibody-specific requirements. Importantly, BD residues essential for binding of nAbs or native folding^30^ were highly conserved across genotypes (Figures 1C and S2), underscoring the value of BD as a promising site of vulnerability for pan-genotypic HCV vaccine design.

### BD nAbs derived from diverse lineages exhibit convergent CDRH3 motifs

In contrast to AR3-targeting nAbs, which are predominantly restricted to the heavy chain variable gene germline *IGHV1-69*, BD nAbs exhibit greater germline diversity (Figure 2A and Table S1). Among the 25 BD nAbs analyzed, 13 (52%) originated from IGHV1 gene family, including 6 (24%) from *IGHV1-18,* and 5 (20%) from *IGHV1-69*. Additional antibodies were derived from the IGHV3 and IGHV4 families. Most antibodies exhibited moderate levels (4%-18%) of somatic hypermutation (SHM) and featured average to longer (13-25 residues) CDRH3 loops (Figure 2B). IGHD3 gene fragments, particularly *IGHD3-10* (20%) and *IGHD3-3* (16%), were most frequently employed, with IGHJ4 (52%) as the dominant joining segment (Figure 2C).

**Figure 2.**
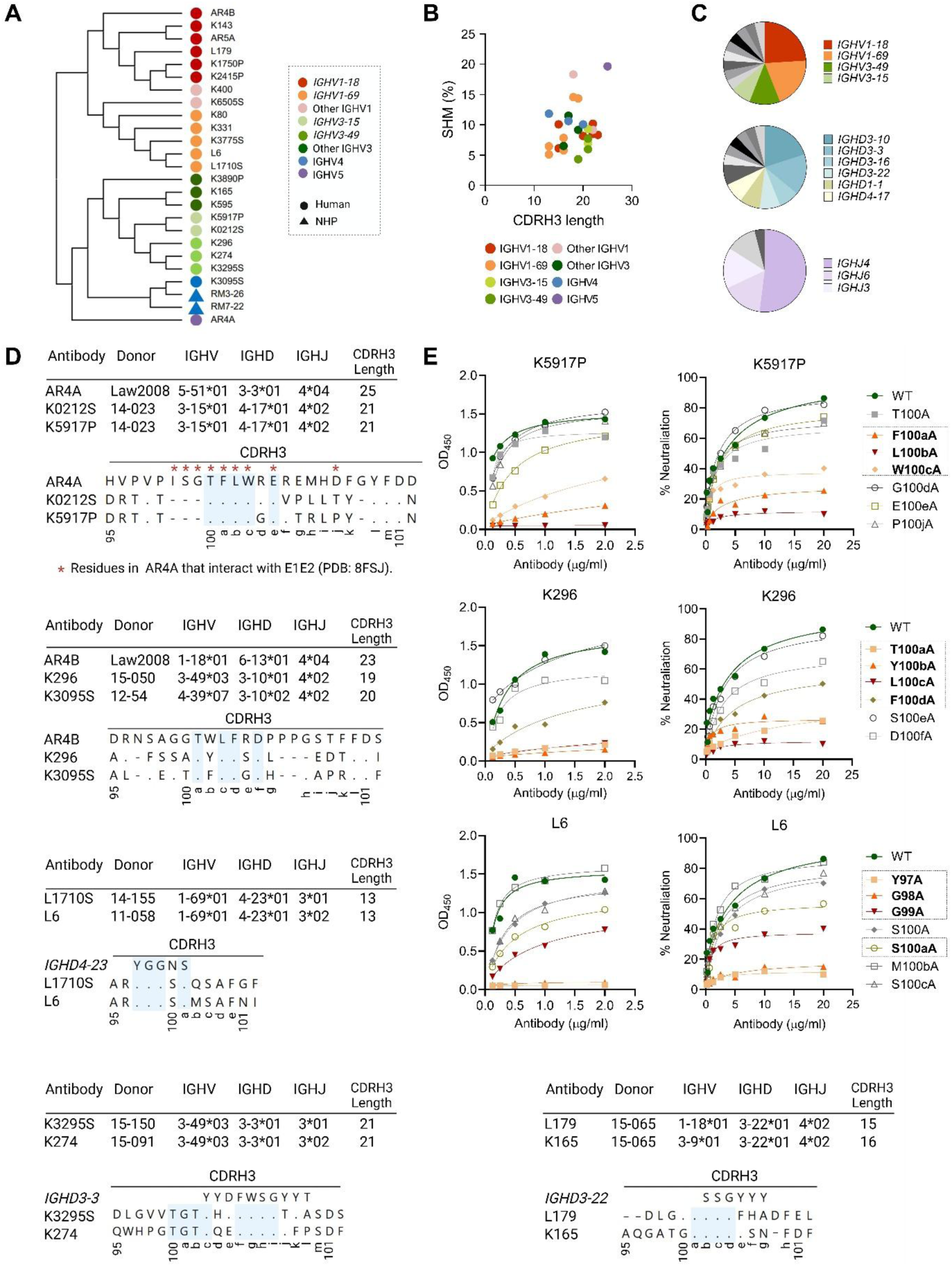
Sequencing characteristics of HCV BD nAbs. (A) Maximum likelihood (ML) phylogenetic tree depicting relationships among heavy-chain amino acid sequences of BD nAbs (n = 25) isolated from humans and NHPs after infection or immunization. (B) SHM frequency and CDRH3 length (Kabat numbering) of BD nAbs. Each dot represents an individual antibody. (C) Germline gene usage distribution of IGHV, IGHD, and IGHJ among BD nAbs. (D) Sequence motifs in CDRH3 (highlighted in light blue) shared across BD nAbs. Asterisks mark epitope-contacting residues in AR4A bound to sE1E2.SZ (PDB: 8FSJ). Kabat numbering is shown below the aligned antibody sequences. (E) ELISA binding and HCVpp neutralization against H77 stain by antibody variants containing defined mutations within CDRH3.

Given the well-established importance of CDRH3 in mediating antigen recognition, we analyzed the CDRH3 characteristics of BD nAbs. Despite originating from diverse germline genes and donors, some antibodies converged on highly similar CDRH3 motifs (Figures 2D and Table S1), suggesting common structural solutions for epitope recognition. For instance, antibodies K0212S and K5917P (both encoded by *IGHV3-15*01*, *IGHD4-17*01* and *IGHJ4*02*) share a conserved TFLWxE motif with AR4A (encoded by *IGHV5-51*01*, *IGHD3-3*01* and *IGHJ4*04*) within their CDRH3 (Figure 2D, upper panel). This motif has previously been identified as a key determinant of AR4A recognition, mediating predominantly hydrophobic interactions with a shallow cleft on E1E2 BD.^12,20^ To evaluate its role in other antibodies, a series of K5917P heavy-chain variants were constructed, each carrying a single alanine substitution in the motif, while maintaining the native light chain pairing. Functional analyses showed that single-residue substitutions at F100a, L100b, and W100c led to substantial reductions in both antigen binding and neutralization activity (Figure 2E, upper panel), underscoring the critical role of this motif in epitope recognition. Similarly, antibodies K296 (encoded by *IGHV3-49*03*, *IGHD3-10*01* and *IGHJ4*02*), K3095S (encoded by *IGHV4-39*07*, *IGHD3-10*02* and *IGHJ4*02*) and AR4B (encoded by *IGHV1-18*01*, *IGHD6-13*01* and *IGHJ4*04*) share a TxLFxD motif within their CDRH3 (Figure 2D, 2nd panel). Alanine substitutions at T100a, Y100b, L100c, and F100d in K296 each resulted in >50% reduction in binding and neutralization (Figure 2E, middle panel).

Beyond these examples, other antibodies exhibited shared IGHD usage with convergent CDRH3 motifs. For example, L1710S and L6, both encoded by *IGHV1-69*01*, *IGHD4-23*01*, and *IGHJ3* and isolated from different donors, shared a YGGSS motif derived from *IGHD4-23* (Figure 2D, 3rd panel). Alanine substitutions at Y97, G98, G99 or S100a in L6 greatly reduced binding and neutralization, highlighting the structural importance of this motif (Figure 2E, lower panel). Likewise, K3295S/K274 and L179/K165 featured TGTYxxFWSG and DSSG motifs derived from *IGHD3-3* and *IGHD3-22*, respectively (Figure 2D, lower panels). These patterns highlight recurrent D gene-driven recombination contributing to BD recognition.

BD nAbs predominantly utilized kappa light chains, most frequently encoded by *IGKV1-5* and *IGKV1-39*, with *IGKJ1* and *IGKJ4* as the most common J gene segments (Figure S3A). Unlike AR3-specific nAbs, whose antigen recognition is largely heavy chain-dependent,^18,31,32^ BD nAbs exhibited low to moderate dependence on their light chains, as evidenced by the marked reduction in binding activity upon substitution with unrelated light chains (Figure S3B). Collectively, these findings demonstrate that BD nAbs possess distinct genetic features compared with AR3-targeting nAbs, originating from a broad range of B cell lineages. Despite this diversity, they reproducibly converged on public CDRH3 motifs, employing common structural strategies for BD recognition among unrelated clones.

### BD bnAbs recognize overlapping epitopes across AR4, AR5 and back layer

We recently reported the epitope mapping of AR4A and AR5A using nsEM.^26^ Following a systematic workflow (Figure S4), we generated 2D class averages and 3D reconstructions from complexes of antigen-binding fragments (Fab) with a His-tagged soluble E1E2 (sE1E2) antigen derived from the HCV-1 strain, in which sE1E2 was fused to a truncated SpyTag/SpyCatcher (SPYΔN), termed HCV-1 sE1E2.Cut_1+2_.SPYΔN-His_6_ (or HCV-1 SpyE1E2; Table S2). The HCV-1 SpyE1E2 antigen adopted a native-like confirmation consistent with recent cryo-EM structures.^12,20,26^ These analyses revealed that AR4A and AR5A can co-bind E1E2 at contiguous but non-overlapping epitopes.

Applying the same nsEM strategy, we examined 12 newly isolated BD nAbs in complex with HCV-1 SpyE1E2 and reference antibodies AR3C and/or AR1B. The reconstructions showed that BD-targeting antibodies bound overlapping yet distinct epitopes centered on AR4 and AR5, with variable extensions into the E2 back layer (Figures 3A and 3B). For example, antibodies L6, K296, K274, K3775S, RM3-26, and K400 were positioned primarily over AR4, whereas K165, K595, K80, and K331 recognized epitopes spanning both AR4 and AR5. Notably, RM3-26, K400, L179, K331, and L1710S exhibited increased contacts with the AR2 back layer region. These patterns were consistent with results from competition ELISA and alanine-scanning mutagenesis (Figures 1A and 1B). Together, these data define new epitopes targeted by BD nAbs, including those bridging the AR4/AR5 interface and others recognizing both the BD and back layer. These findings indicate that BD bnAbs converge on a common antigenic surface but engage it through distinct binding modes, revealing multiple structural solutions to BD-directed neutralization.

**Figure 3.**
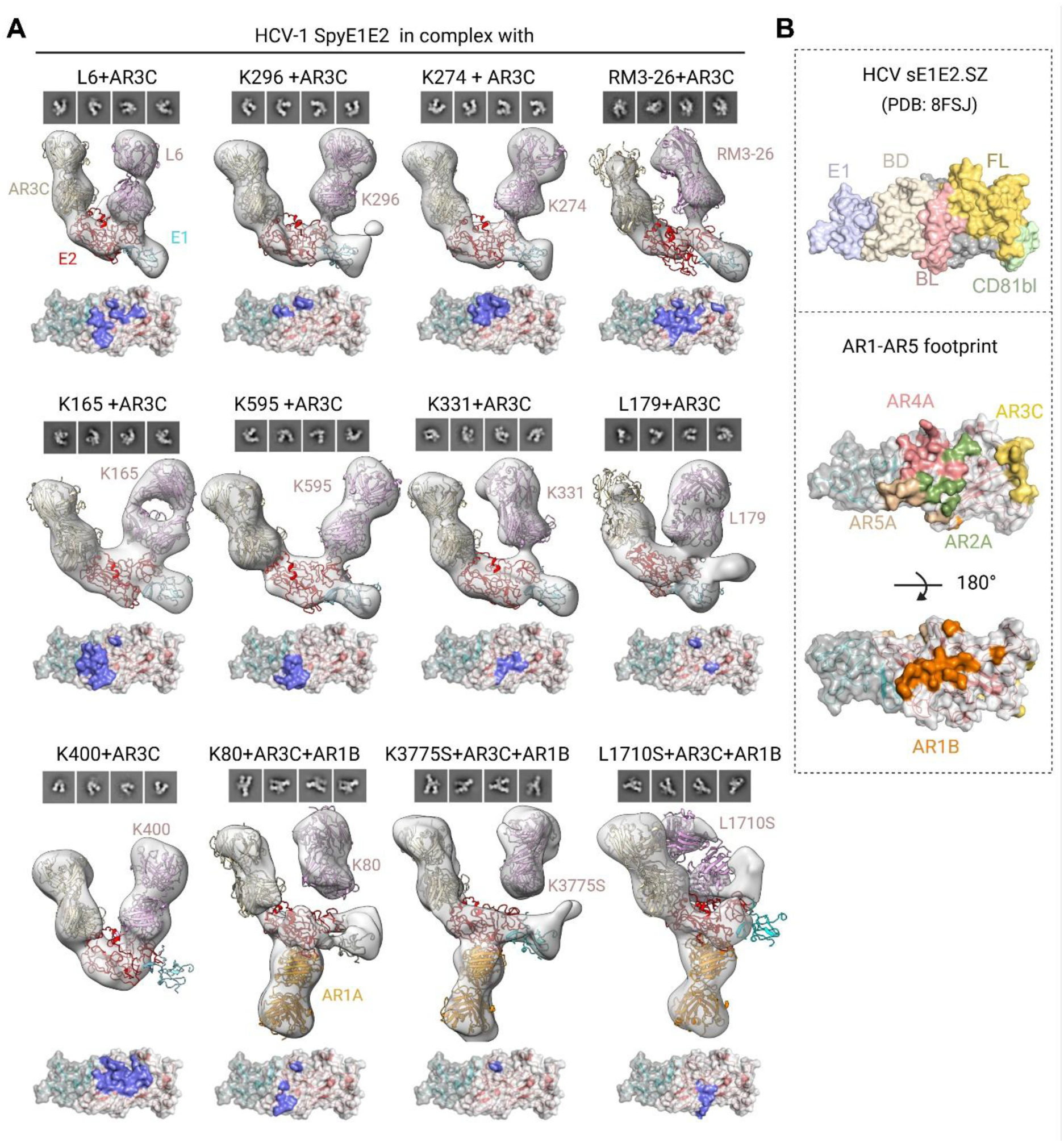
Epitope mapping of BD nAbs by nsEM. (A) 2D classification representatives, 3D reconstructed model and epitope footprint analysis for HCV-1 SpyE1E2 in complex with Fab BD nAbs, with AR3C and/or AR1B as reference antibodies. 3D reconstructed map was shown in surfaces. The Fab structural models were generated by SWISS modeling,^61^ except K595 and K400. K595 and K400 structures were obtained by X-ray crystallography (Table S3). The footprints on the surfaces are shown using the cryo-EM structure of sE1E2.SZ (PDB: 8FSJ). (B) Upper panel: Surface representation subregions of 1b09 sE1E2.SZ (PDB: 8FSJ), including E1, BD, BL (BL), front layer (FL), CD81-binding loop (CD81bl), and the rest of E1E2 are colored in lavender, ivory wheat, light pink, yellow orange, praseodymium, gray, respectively. Lower panel: Epitope footprints of reference antibodies AR1B, AR2A, AR3C, AR4A, and AR5A mapped on sE1E2.SZ.

### Crystal structure of RM3-26 in complex with H77 E2c3

We next sought to determine high-resolution X-ray crystal structures of these BD nAbs in complex with a full-length E1E2 (e.g., HCV-1 SypE1E2). However, despite extensive optimization, crystallization trials either failed to produce crystals or produced crystals with limited diffraction. Interestingly, unlike most BD-targeting antibodies that completely lost binding to soluble E2 constructs, a few BD nAbs retained weak but specific affinity relative to the native E1E2 complex. K400, for instance, bound both H77 and HK6a E2c3 (an E2 core domain variant comprising residues 412–459, 486–568, and 598–645; Table S2) with a dissociation constant (Kd) of ∼20 nM and HCV-1 E2ecto with a Kd of 108 nM (Figure 1B). Another antibody, RM3-26, showed no detectable binding to HCV-1 E2ecto (Figure 1A). However, its Fab displayed very weak binding to H77 E2c3 (Kd = 305 nM) but no detectable interaction with HK6a E2c3 or HCV-1 E2ecto (Figure S1B). We therefore attempted co-crystallization of K400 and RM3-26 with E2c3 constructs. While crystals of K400 contained only Fab molecules (Table S3), likely due to rapid dissociation from E2c3 (Figure S1B), the structure of RM3-26 in complex with H77 E2c3 was successfully determined at 2.24 Å resolution (Table S3).

In the RM3-26-E2c3 complex structure, the RM3-26 Fab was clearly resolved (Figure 4A). In contrast, H77 E2c3 exhibited weaker density overall, with the most defined density corresponding to the back layer (residues 615–645). This region formed the principal binding interface with RM3-26, consistent with the epitope mapping by alanine scanning and nsEM (Figures 1C and 3A). The RM3-26-E2c3 interaction is almost entirely mediated by the antibody heavy chain, which accounts for 99% of the buried surface area (BSA; 494 Å² out of a total 511 Å²) (Figures 4B-4D). This observation is in line with functional data showing that RM3-26 binding was only slightly reduced upon light chain exchange (Figure S3B). The majority of contacts are with CDRH1 (BSA of 197 Å²) and CDRH3 (BSA of 217 Å²), which primarily engage the E2 back layer at residues 614, 615, 622-629, and 641-645 (Figures 4B-4D). Several of these residues, including L615, N623, T625, and F627, are highly conserved across HCV genotypes (Figure S2).

**Figure 4.**
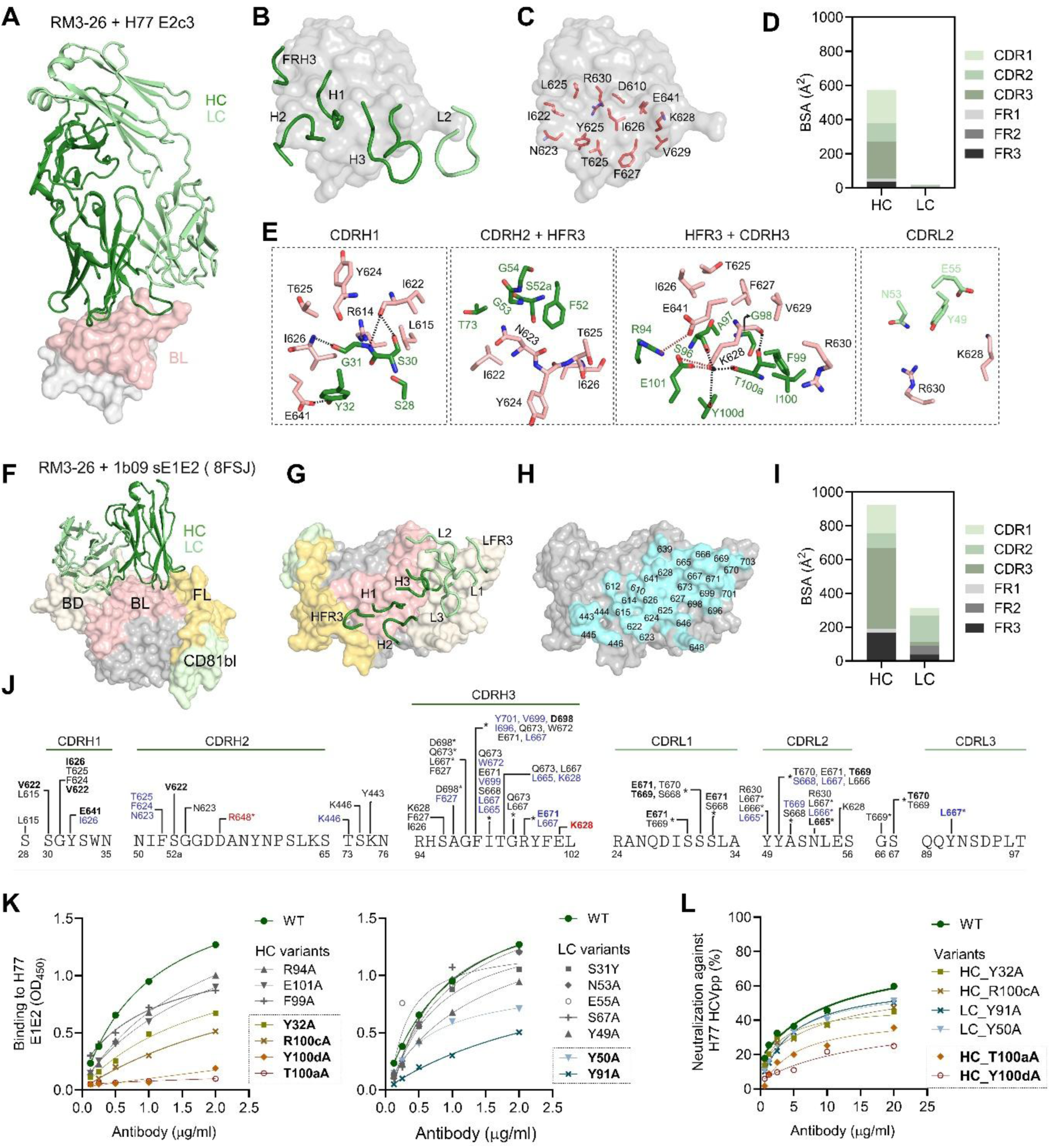
Crystal structure of RM3-26 in complex with H77 E2c3. (A) The overall complex structure of RM3-26 and H77 E2c3. RM3-26 is shown in a ribbon representation with HC (dark green) and LC (pale green). H77 E2c3 is shown as a light gray solid surface with the back layer (BL) colored pink. (B) Position of RM3-26 CDRH, CDRL, HFR on H77 E2c3 (gray surface). The CDRH, CDRL, HFR are shown as tubes. (C) The epitope of RM3-26 on H77 E2c3 gray surface. The epitope residues of H77 E2c3 that have BSA > 0 Å^2^ are shown in salmon. (D) Buried surface area (BSA) of RM3-26 HC and LC subregions on H77 E2c3. (E) Detailed binding interactions between RM3-26 HC (dark green), LC (pale green), and H77 E2c3 BL (salmon). The epitope and paratope residues are labeled. The hydrogen bonds and ionic interactions are shown in dotted black and red lines, respectively. (F) Predicted model of RM3-26 bound to 1b09 sE1E2.SZ (PDB: 8FSJ). The RM3-26+H77 E2c3 complex was superimposed onto the sE1E2.SZ structure (PDB: 8FSJ) based on the E2 region to generate the full-length complex model. RM3-26 HC (dark green) and LC (pale green) are shown as cartoons. sE1E2.SZ subregions are displayed as surface representations, colored as in Figure 3B. (G) Modeled position of RM3-26 CDRH, CDRL, and FR regions (shown as ribbons) on the sE1E2.SZ surface. (H) Predicted epitopes (cyan) of RM3-26 on sE1E2.SZ (gray) surface. The residues of E2 that are predicted to contact RM3-26 are indicated. (I) BSA of RM3-26 HC and LC subregions on the model of sE1E2.SZ. (J) Schematic overview of the interactions between E2 and RM3-26. Hydrophobic (blue), hydrogen-bonding (bold), and ionic (red) residues are indicated. Asterisks (*) denote predicted contact residues derived from the 1b09 sE1E2.SZ model with RM3-26. (I) ELISA binding (left) and HCVpp neutralization (right) of RM3-26 antibody variants carrying alanine substitutions at key heavy-chain contact residues against the H77 viral strain. (K and L) Functional evaluation of RM3-26 variants versus the wildtype (WT) antibody by binding ELISA (K) and HCVpp neutralization (L). Each HC variant was paired with WT LC, and vice versa.

CDRH1 of RM3-26 forms multiple hydrogen bonds with the E2 back layer, including S30 and G31 with I622, G31 with I626 main chain, and Y32 with E641, along with a hydrophobic contact with I626 (Figure 4E and Tables S4 and S5). In CDRH2, F52 makes hydrophobic interactions with N623 and T625 (Figure 4E and Tables S3 and S4). CDRH3 mediates extensive polar and electrostatic contacts, including salt bridges between R94 and E641 and between E101 and K628 (Figure 4E and Table S4). A hydrogen-bonding cluster formed by S96, T100a, Y100d, and E101 engages the K628 side chain, with T100a also hydrogen bonding with the K628 main chain and forming a hydrophobic interaction with the same residue (Figure 4E and Table S5). The light chain contributes modestly through CDRL2 residues Y49, N53, and E55, which are buried in the interface between K628 and R630 of the E2 back layer (Figure 4E).

### Structural model of RM3-26 on E1E2

The RM3-26/E2c3 interface is relatively small compared to classical Fab-antigen complexes (500 Å² vs. 989 Å^2^ for Fab AR3C/H77 E2 core^31^ and 834 Å^2^ for AR3A/HK6a E2c3.^33^ However, structural superposition of the RM3-26/E2c3 complex onto the sE1E2.SZ structure (PDB: 8FSJ) suggests that the RM3-26 epitope would be substantially larger in the context of full-length E1E2, with an estimated BSA of ∼1,070 Å² (Figures 4F-4I).

In the model for RM3-26 bound to E1E2, the RM3-26 interactions were also dominated by the heavy chain at the E2 BL, with auxiliary contacts to the E2 BD (Figure 4G-4I). These contacts include residues from the heavy-chain CDRH3 and the light-chain CDRL1–3, particularly LFR3 (Figure 4G and 4J). Predicted hydrogen bonds formed between F99-D698 and R100c-E671 in RM3-26 CDRH3, together with contributions from S31, Y50/N53, S67, and Y91 from RM3-26 light chain, suggesting a coordinated recognition of the BD (Figure 4J). This structural model is further supported by nsEM and alanine scanning, which confirm BD recognition and dependence on key BD residues (Figures 1B and 3A).

To pinpoint the contribution of RM3-26 residues, individual alanine substitutions were introduced at key contact positions. Substitutions at T100a or Y100d in RM3-26 CDRH3 abolished both binding and neutralization, underscoring their essential role in antigen recognition (Figure 4K, 4L). Mutations at CDRH1 Y32, CDRH3 R100c, CDRL2 Y50, and CDRL3 Y91 result in moderate binding reductions (Figure 4J). Collectively, the crystal structure of RM3-26 in complex with H77 E2c3 provides a clear visualization of the BL region of its epitope. Structural superposition RM3-26 onto E1E2, together with epitope and paratope mapping, confirmed that RM3-26 recognizes an epitope spanning the BL and BD on E1E2.

### RM3-26 employs a binding mode similar to that of hcab40 to recognize an epitope on the E2 back layer

We compared the crystal structure of the RM3-26-E2c3 complex with previously determined complex structures of the human back layer-targeting nAb hcab40 bound to E2ecto (PDB: 8W0X),^34^ the mouse back layer-targeting non-nAb 2A12 bound to E2ecto + BD (residues 646-712, PDB: 8DK6),^35^ the human BD-targeting bnAb AR4A bound to sE1E2.SZ (PDB: 8FSJ),^20^ and the human domain A/back layer-targeting non-neutralizing antibody CBH4B bound to E2ecto (PDB: 8TXQ).^36^ Despite differences in sequence, species origin, and epitope specificity, these antibodies shared a common feature of engaging the E2 back layer. RM3-26, Hcab40, and AR4A exhibited similar overall binding modes in their recognition of the E2 back layer, which differed markedly from those of AR3-directed antibodies such as human bnAb HEPC74 (Figure 5A).

**Figure 5.**
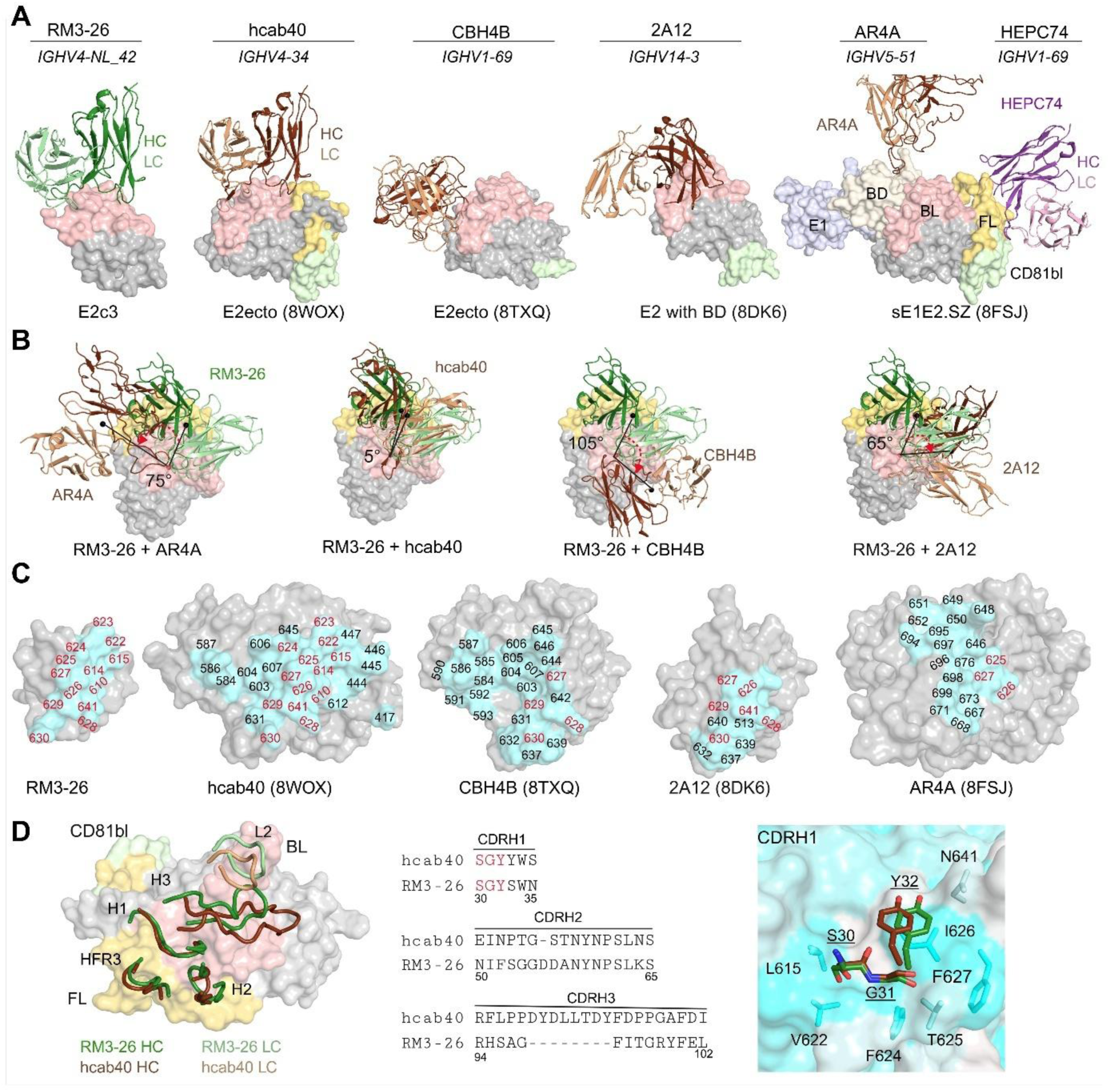
RM3-26 targets the AR4A epitope with an approach angle similar to that of human hcab40. (A) Superposition of E2 from RM3-26, hcab40 (PDB: 8WOX), CBH4B (PDB: 8TXQ), 2A12 (PDB: 8DK6), AR4A, and HEPC74 (PDB: 8FSJ) complexes show that RM3-26, hcab40, CBH4B, and AR4A share similar back-layer binding modes, distinct from HEPC74 targeting the AR3 region. Fabs are shown as cartoons with E2 in surface representation. HC and LC of RM3-26 are dark green and pale green; HEPC74, purple and light pink; others, dark and light brown. sE1E2.SZ subregions are colored as in Figure 3B. (B) The binding angle of RM3-26 with other back layer-targeting antibodies. The HC and LC of Fab and E2 are same as A. (C) The epitopes of BL-targeting antibodies are highlighted on the E2 surfaces in cyan. Residues (red labels) that overlap with the RM3-26 epitope are located within the BL region of E2. The E2 surface is shown in gray. (D) RM3-26 and human hcab40 exhibit similar binding modes to E2. Left, the CDRH1–3, HFR3, and L2 regions of RM3-26 and hcab40 adopt similar orientations on the E2 surface. Middle, sequence alignment of the CDRH1-3 regions of RM3-26 and hcab40. The shared S^30^G^31^Y^32^ motif in CDRH1 is highlighted in red. Right, structural overlay showing that the CDRH1 of RM3-26 and hcab40 encode the S^30^G^31^Y^32^ motif and engage E2 in a similar binding mode. This motif is positioned within a conserved hydrophobic pocket. E2 residues are shown in cyan, while RM3-26 and hcab40 residues are colored dark green and brown, respectively. The E2 surface is colored according to hydrophobicity, with cyan representing hydrophobic regions and white representing hydrophilic regions.

AR4A has a different approach angle of about 75° anti-clockwise away from the RM3-26 antibody (Figure 5B), however, the approach angle of RM3-26 is similar to that of human hcab40 with an approximate difference of 5°. By contrast, CBH4B and 2A12 are 105° and 65° clockwise away from the RM3-26 antibody, respectively (Figure 5B). All of these antibodies recognize epitopes containing back layer residues (Figure 5C).

Comparison of the epitopes recognized by these antibodies revealed that RM3-26 engages overlapping regions within the E2 back layer: residues 627-630 with CBH4B, 627-630 and 641 with 2A12, and 625-627 with AR4A (Figure 5C). Among these, RM3-26 and hcab40 recognize nearly identical epitopes within the E2 back layer, with shared contacts at residues 615, 622-628 and 641 (Figure 5C and Table S4). Notably, F627 is a key contact residue and is broadly conserved across multiple HCV genotypes (Figures 5C and S2).

Strikingly, structural comparison revealed that the CDRH1-3, HFR3, and CDRL2 of RM3-26 and human hcab40 are similarly positioned (Figure 5D). Both antibodies contain an SGY motif within CDRH1 that mediates hydrogen bonding: S30 and G31 contact E2 residues I622V and the main chain of I626, respectively (Figure 5D and Table S5). Their SGY motifs are surrounded by a hydrophobic pocket composed of conserved hydrophobic residues L615, V622, F624, T625, I626, F627, and N641 (Figures 5D right panel and S2; Table S5). In addition, K628 hydrogen bonds with RM3-26 residues S96, T100a, Y100d, and E101, or with hcab40 P98 and Y100 (Table S5). These structural and sequence parallels underscore the convergent binding strategies of macaque RM3-26 and human hcab40 despite their distinct origins.

When each antibody was superimposed onto native E1E2 structure containing the transmembrane region (full E1E2; PDB: 8RJJ) (Figure S5A and Table S2), the long CDRH3 of hcab40 and the entire heavy chain of CBH4B sterically clashed with the core of the E2 BD, whereas neither RM3-26 nor AR4A exhibited such clashes (Figure S5B). In addition, CBH4B and 2A12 also collided with E1 (Figure S5). The CDRH3 of hcab40 adopts a straight elongated hairpin conformation, while the CDRH3 loops of RM3-26 and AR4A are more bent, thereby avoiding steric conflict with the E2 BD (Figure S5). nsEM confirms that nAb K400 binds a similar epitope as RM3-26 (Figure 3) and our unliganded crystal structure of Fab K400 (Figure S6 and Table S3) shows that its CDRH3 also adopts a bent hairpin conformation.

### BD nAbs achieve affinity maturation and broad neutralization with minimal SHM at contact residues

To delineate the molecular mechanisms underlying BD-directed antibody maturation, we investigated two structurally defined representatives: RM3-26 and AR4A. RM3-26 is a nAb with moderate breadth and potency (Figure 1A), isolated from an NHP immunized with HCV-1 E1E2.^1^ Its heavy chain (derived from *IGHV4-NL_42*01_S3666*, *IGHD1-2*01*, and *IGHJ1-1*01*) contains 13 SHMs at the amino acid level, of which four (Y52F in CDRH2, and S53G, V98G and V99F in CDRH3) directly contact HCV E2 (Figures 6A and 6B). The light chain (originates from *IGKV3-20*01* and *IGKJ5*01*) carries 9 SHMs, none of which are located at the antibody-antigen interface. To define the minimal requirements for RM3-26 maturation and function, we generated a panel of heavy and light chain germline revertants and partially matured variants, including the germline precursor (GL), GL with mature CDR3 (GM), and GM derivatives harboring key SHMs (Figure 6A). Each heavy chain variant was expressed with the wild-type light chain, and vice versa, for functional evaluation. The RM3-26 heavy chain GL variant exhibited minimal activity, whereas GM restored partial function against several viral strains (Figure 6C). A single SHM at position 52, 52a, or 53 in the GM background significantly enhanced antibody function, and combinations of SHMs at 52 and 53 or incorporation of the mature CDRH2 further improved function (Figure 6C). Notably, pairing the RM3-26 wild-type heavy chain with a germline light chain retained near-complete activity (Figure 6C), consistent with a dominant role of the heavy chain in affinity maturation.

**Figure 6.**
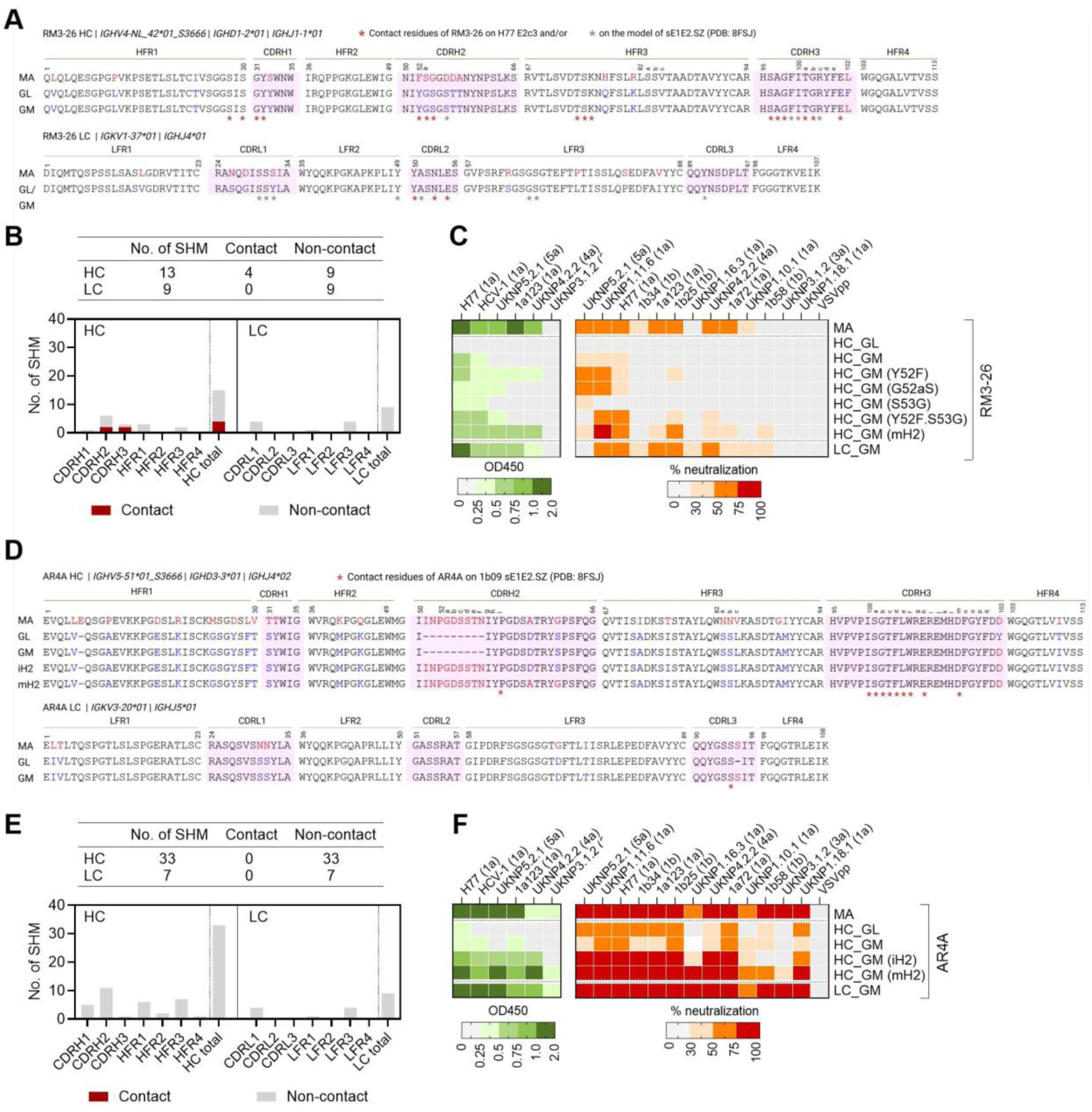
Minimal SHM requirements for functional maturation of BD nAbs RM3-26 and AR4A. (A) Sequence alignment of RM3-26 heavy (HC) and light (LC) chain inferred germline precursor (GL), GL with mature CDR3 (GM), and mature antibody (MA). SHMs in MA are highlighted in red, corresponding germline residues in blue, and E2-contacting residues are denoted by an asterisk (*). Germline gene assignments and Kabat numbering are annotated. Each HC variant was paired with the wild-type LC, and vice versa, for functional evaluation. (B) Number and distribution of SHMs in the HC and LC of the RM3-26 antibody. (C) Cross-binding and cross-neutralization of RM3-26 variants against representative tier 1-4 HCV strains. (D) Sequence alignment of AR4A HC and LC MA, GL and GM, as HC GM variants containing either an insertion in CDRH2 (iH2) or the mature CDRH2 (mH2). SHMs in MA are colored red, corresponding germline residues are blue, and contact residues are denoted by an asterisk (*). Germline gene assignments and Kabat numbering are annotated. (E) Number and distribution of SHMs in the HC and LC of the AR4A antibody. (F) Cross-binding and cross-neutralization of AR4A variants against representative tier 1-4 HCV strains.

AR4A represents one of the most potent and broadly neutralizing HCV bnAbs identified to date.^37^ It was isolated from a chronically infected individual.^21^ The heavy chain (derived from *IGHV5-51*, *IGHD3-16*02*, and *IGHJ4*02*) accumulated 33 SHMs, while the light chain (encoded by *IGKV1-37*01* and *IGHJ4*01*) acquired 7 (Figures 6D and 6D). Strikingly, most of these mutations occur within framework regions, and none directly contribute to antigen recognition. Although AR4A primarily engages E1E2 through its CDRH3 loop (accounts for 89% of BSA),^37^ its heavy chain GL and GM variants (both retaining all contact residues in CDRH3) exhibited only weak binding and limited neutralization activity (Figure 6F). In addition, AR4A heavy chain features a unique 9-amino acid insertion and two SHMs in CDRH2, while the putative contact residue Y52i^20^ is retained from the germline (Figure 6D). Inclusion of this insertion or the matured CDRH2 greatly improved binding and neutralization (Figure 6F), indicating that non-contact SHMs are critical for full functional maturation. Similar to RM3-26, pairing the AR4A wild-type heavy chain with a germline light chain had little impact on binding or neutralization (Figure 6F). Collectively, these data reveal two distinct maturation strategies for RM3-26 and AR4A: RM3-26 relies primarily on targeted SHMs within its contact residues to achieve function, whereas AR4A requires extensive non-contact SHMs that reinforce structural stability and optimize paratope geometry to enable exceptional breadth.

### Combination of BD and NF bnAbs enhanced neutralizing breadth and potency

Despite the broad neutralizing activity of some HCV bnAbs (e.g., AR4A and AR3C), it is unlikely that a single bnAb, or a bnAb lineage, will provide coverage against all circulating strains. Thus, inducing bnAbs that target multiple antigenic sites offers a promising approach to prevent viral escape. NF- and BD-targeting bnAbs constitute two of the most potent and broadly neutralizing classes identified to date. They differ in their epitopes, sequence usage, and structural recognition properties (Figure 7A and Table 1). Because they recognize distinct antigenic regions and neutralize by different mechanisms, BD and NF bnAbs are promising partners for vaccine design aimed at eliciting synergistic protective responses.

**Figure 7.**
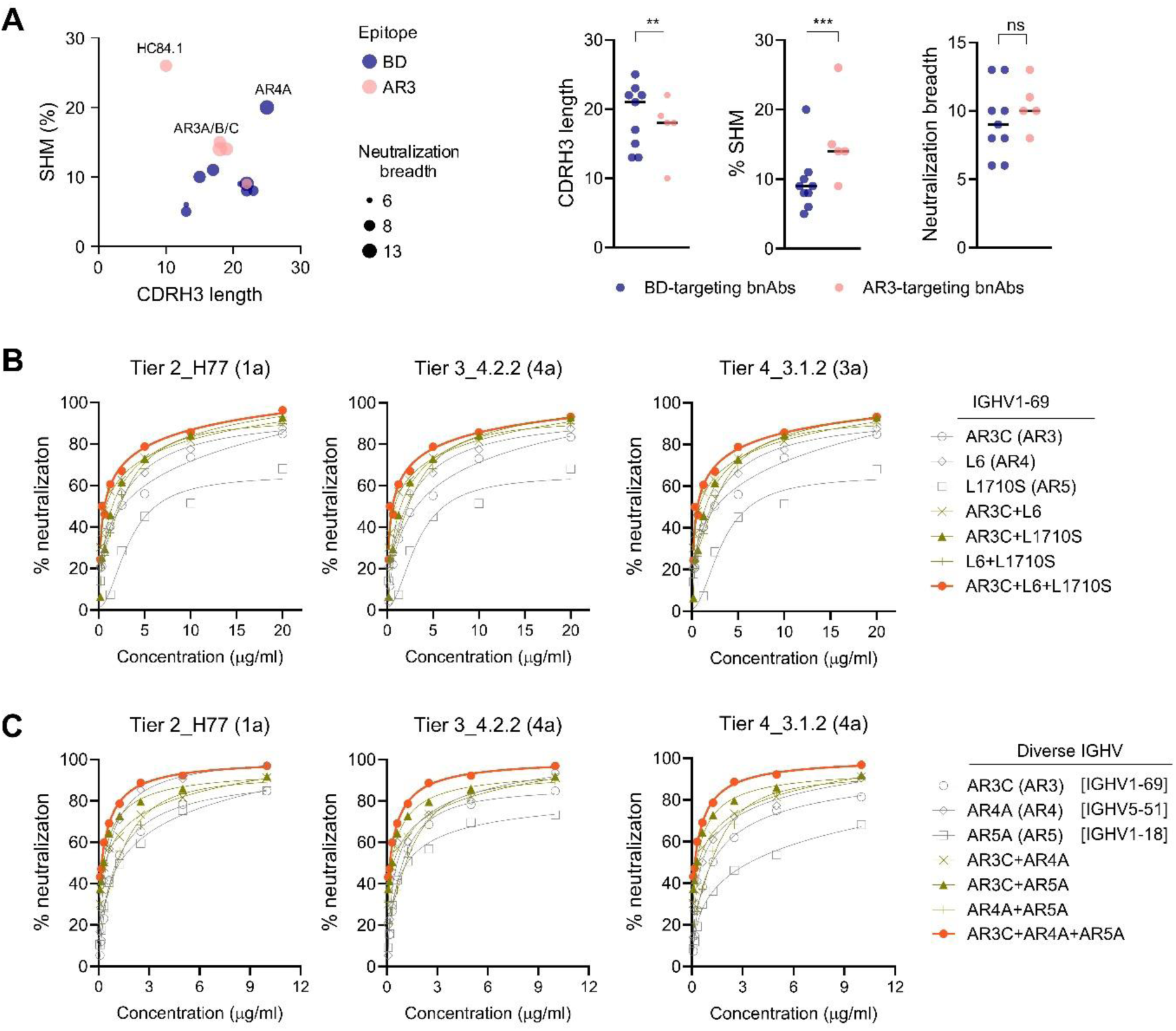
Cooperative effects of NF and BD bnAbs in neutralization. (A) SHM, CDRH3 length and neutralization breadth of representative AR3 and BD bnAbs. (B) HCVpp neutralization against H77, UKNP4.2.2, and UKNP3.1.2 strains by combinations of NF- and BD-specific nAbs encoded by *IGHV1-69*. (C) HCVpp neutralization of H77, UKNP4.2.2, and UKNP3.1.2 strains by combinations of AR3- and BD-specific nAbs utilizing diverse IGHV gene segments.

**Table 1.**
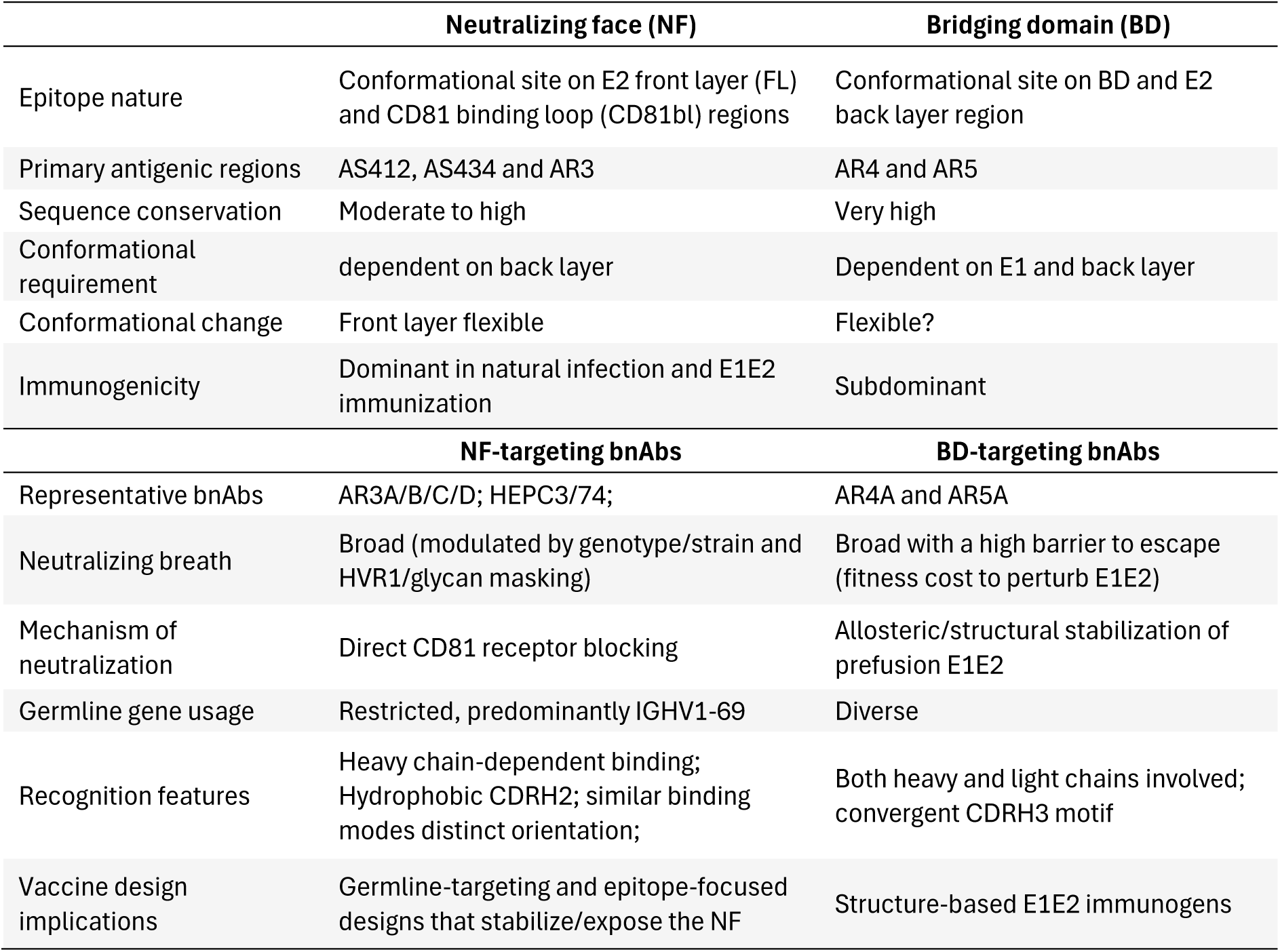
Comparison of NF and BD epitope and bnAbs.

To evaluate their cooperative potential, we tested combinations of BD- and NF-targeting bnAbs derived from both shared and diverse lineages. Within the *IGHV1-69*-encoded group, AR3C (targeting AR3), L6 (targeting AR4) and L1710S (targeting AR5) each displayed moderate to broad neutralization breadth individually. However, their pairwise or triple combinations markedly enhanced potency and breadth, with additivity observed against multiple tier 2-4 neutralization-resistant viral strains (Figure 7B). A similar pattern was observed with bnAbs derived from distinct germlines: AR3C (encoded by *IGHV1-69*), AR4A (encoded by *IGHV5-51*), and AR5A (encoded by *IGHV1-18*), where combinations, particularly the triple, achieved greater neutralization potency and breadth (Figure 7C). Together, these findings demonstrate that AR3- and BD-targeting bnAbs act either additively or synergistically, highlighting their complementarity and underscoring the value of vaccines designed to elicit multi-epitope bnAb responses.

## Discussion

Eliciting bnAbs capable of potently neutralizing a broad spectrum of viral strains remains a major challenge for HCV vaccine development. Both NF and BD on the E1E2 complex are attractive templates, but BD epitopes and their cognate bnAbs are still poorly understood. Here, by characterizing a panel of 25 BD-specific nAbs isolated from infection and immunization, we revealed their molecular and structural determinants, offering new insights into this underexplored antibody class to inform strategies for rational vaccine design.

BD nAbs differ greatly from AR3 nAbs in their epitopes, developmental origins, and structural/functional properties (Table 1). They can arise from diverse B cell lineages, and distinct antibody lineages can converge on shared CDRH3 features (Figure 2). This convergence implies that the BD imposes stringent structural constraints on antibody recognition, selecting recurrent sequence motifs and paratope architectures. Such patterns parallel public antibody solutions observed against conserved HCV E1E2 epitopes, including those on the E2 NF. Notably, *IGHV1-69* is frequently utilized by both BD and AR3 bnAbs, although it is less dominant among BD-specific antibodies (Figure 2C). This observation suggests partial convergence on a germline-encoded pathway. Characterizing the molecular features of *IGHV1-69*-derived bnAbs targeting both antigenic sites could support the development of germline-targeting vaccine strategies that reproducibly prime appropriate precursors, enabling the coordinated elicitation of antibodies to multiple conserved epitopes and improving overall neutralization breadth.

AR4 and AR5 represent the two major non-overlapping antigenic sites within the BD.^26^ Detailed epitope mapping of these newly identified BD nAbs by competition ELISA, alanine scanning and nsEM revealed a subset (e.g., K165 and K595) targeting composite epitopes spanning AR4 and AR5 (Figures 1A, 1B and 3A). This finding indicates that the BD likely forms a quaternary, composite interface consisting of a continuum of overlapping epitopes rather than a single discrete site. In addition, some BD nAbs (e.g., RM3-26 and K400) also engage the E2 back layer, which enabled structural analysis of these antibodies in complex with E2 alone, given the substantial challenges of crystallizing BD nAbs with the full E1E2 heterodimer.

To date, no X-ray crystal structure of a BD-targeting antibody bound to E1E2 has been reported. The structure of AR4A in complex with E1E2 was recently determined by cryo-EM, representing the only available high-resolution view of a BD epitope on native-like E1E2.^12,20^ Despite extensive efforts, we were unable to obtain a BD antibody-E1E2 co-crystallization. Nevertheless, we determined a high-resolution crystal structure of a nonhuman primate BD antibody, RM3-26, in complex with the E2c3, revealing its epitope extending to the back layer (Figure 4). The binding orientation of RM3-26 closely mirrors that of the human back layer-targeting antibody hcab40 (Figure 5), highlighting conserved epitope recognition across species.

Furthermore, superposition of E2 back layer-targeting antibodies, RM3-26, hcab40, CBH4B, and AR4A, onto full E1E2 (PDB: 8RJJ) positioned these antibodies near the ectodomain-membrane junction (Figure S5). This localization suggests that such antibodies may inhibit membrane fusion by locking or destabilizing prefusion intermediates, thereby preventing insertion of the fusion peptide (within the BD or helical segment of E2 adjacent to the TM region) into the target membrane. This mechanism may parallel those of MPER-directed antibodies against HIV (e.g., 10E8) and fusion-peptide–targeting antibodies against influenza. Some viral glycoproteins, such as HIV gp41, Ebola GP, SARS-CoV-2 S2, and influenza HA, undergo major structural rearrangements to facilitate membrane fusion, although this process remains unknown for HCV. Alternatively, the proximity of these epitopes to the viral membrane may enable steric hindrance, with bound antibodies acting as physical barriers that block the close approach of viral and host membranes required for fusion.

In our previous NHP immunization study, BD nAbs (e.g., RM3-26 and RM7-22) were elicited but were subdominant and showed limited neutralization,^1^ underscoring the need for next-generation immunogens to present the BD quaternary epitope more effectively to the immune system. To delineate maturation dependencies, we generated RM3-26 GL revertants and MA variants and compared them with the human bnAb AR4A. The unmutated/germline precursor of RM3-26 neither bound E1E2 nor neutralized virus. By contrast, pairing the germline with a mature CDRH3, together with targeted single SHMs in CDRH2, produced a stepwise gain in affinity and yielded partial neutralizing activity (Figures 6A-6C). These findings indicate that CDRH3 maturation is the principal driver of RM3-26 function, with additional SHMs further refining the paratope. Conversely, AR4A, despite its extensive overall SHM, did not rely on specific mutations for initial recognition (Figures 6D-6F). Most SHMs were located in the framework regions, along with a unique insertion in CDRH2, but none directly contacted the antigen. These non-contact SHMs may help maintain correct paratope orientation and domain stability, and are critical for the development of neutralization breadth and potency.^38,39^ Together, these findings indicate two distinct maturation solutions to BD recognition: RM3-26 depends on CDRH3 pre-configuration with additional SHMs, whereas AR4A requires extensive SHMs outside of the antibody paratope. While it may be difficult to elicit antibodies with equivalent AR4A SHMs, the isolation of other AR4-targeting bnAbs with normal SHM rates underscore the feasibility of targeting BD and AR4 by vaccination. These insights provide valuable guidance for germline-targeting vaccine design.

For highly diverse viruses like HCV, multiepitope vaccine strategies confer robustness by reducing single-site escape, broadening coverage, and leveraging orthogonal neutralization mechanisms. Despite substantial genetic and structural differences between NF- and BD-targeting bnAbs, their combinations yielded additive or synergistic neutralization (Figure 7), consistent with previous observations.^40,41^ These findings support vaccine approaches that intentionally present both antigenic sites, with the explicit goal of co-maturing AR3-, AR4- and AR5-like lineages to maximize potency and breadth. A deeper understanding of the molecular and structural features of antibody–epitope recognition, particularly the lineage determinants (e.g., *IGHV1-69*) and quaternary constraints at the E1E2 interface, will facilitate lineage-aware and structure-based vaccine designs. In sum, we define the HCV E1E2 BD as a structurally conserved, immunologically tractable target and the diverse bnAbs may share convergent recognition motifs. The molecular and structural knowledge gained provides a framework for rational immunogen design aimed at eliciting cross-protective antibody responses.

## Method Details

### B cell sorting by flow cytometry

Before B cell sorting, the biotinylated H77 sE1E2.SZ protein^42^ (generously provided by Dr. Thomas R. Fuerst) was individually conjugated with a PE streptavidin (ThermoFisher Scientific) or a BV421 streptavidin (BioLegend) at a 4:1 molar ratio, as previously described.^43^ The probes were mixed in 1:1 Brilliant Buffer (BD Bioscience) and FACS buffer (PBS with 2% FBS and 2 mM EDTA) with 5 μM free D-biotin.

To isolate E1E2-specific B cells, cryopreserved PBMCs were thawed and stained with LIVE/DEAD™ Fixable Aqua (ThermoFisher Scientific) in 96-well U-bottom plates to exclude non-viable cells. Following blocking with anti-CD81 (5 μg/mL, BD Biosciences) and TruStain FcX™ (BioLegend), cells were labeled with 200 ng of biotinylated E1E2 per fluorophore in 50 μl at 4°C for 30 min. Surface marker antibodies were then added and incubated for an additional 30 min at 4°C in the dark. E1E2-specific B cells were either bulk-sorted on a MoFlo® Astrios EQ (Beckman Coulter) into 0.04% BSA in PBS for single-cell RNA sequencing (scRNA-seq), or individually collected into 96-well PCR plates containing 4 μL of lysis buffer supplemented with RNaseOUT (Invitrogen) and dithiothreitol for single B cell cloning.

### Production of monoclonal antibodies

Ig heavy and light chain sequences obtained from single-cell RNA sequencing or single B cell cloning were either synthesized by GeneArt Gene Synthesis (ThermoFisher Scientific) or amplified directly, and then cloned into human Igγ1, Igκ and Igλ expression vectors as previously described.^1^ Plasmids containing paired antibody heavy- and light-chain sequences were co-transfected (1:1 ratio) with pAdvantage (Promega) into Expi293 suspension cells using ExpiFectamine™ 293 Transfection Kit (ThermoFisher Scientific) according to the manufacturer’s protocol. Antibody-containing supernatants were harvested 14 days post transfection by centrifugation to pellet cells, filtered through 0.22 µm filters and purified over Protein A-Sepharose 4 Fast Flow (Cytiva) column according to the manufacturer’s protocol.

For the biophysical studies, antibody Fab format was expressed by transient co-transfection of the Fab heavy- and light-chain expression vectors into Expi293F cells using ExpiFectamine (ThermoFisher) according to the manufacturer’s protocol. Affinity chromatography using CaptureSelect CH1-XL affinity matrix (ThermoFisher) was used to purify recombinant Fab from culture supernatant followed by size-exclusion chromatography (SEC) using a Superdex 200 column (GE Healthcare). The purity and integrity of recombinant proteins were confirmed by reducing and nonreducing SDS-PAGE.

### ELISA

E1E2 binding ELISA was performed as previously described.^1^ Briefly, Costar High Binding Half-Area 96-well plates (Corning) were coated overnight at 4 °C with 5 μg/ml of *G. nivalis* lectin (GNL, Vector Laboratories). Following blocking with 4% nonfat milk (Bio-Rad) in PBS containing 0.05% Tween-20 (PBS-T), plates were incubated at room temperature for 1 hour with either batch-diluted cell lysates from 293T cells expressing HCV E1E2. Purified monoclonal antibodies (mAbs) were then added at a final concentration of 1 µg/mL and incubated at room temperature for at least 1 hour. Detection was performed using horseradish peroxidase (HRP)-conjugated goat anti-human IgG Fc-specific antibody (1:2,000; Jackson ImmunoResearch).

For antibody competition analysis, GNL-captured HCV-1 E1E2 was blocked with dilutions of sera/plasma (1:50) or mAbs (20 μg/ml) and incubated for 30 min at room temperature, followed by addition of biotinylated mAb diluted to 75% maximal binding level. Binding was detected using HRP-conjugated streptavidin (1:2,000; Thermo Fisher Scientific). Results are expressed as the percentage of inhibition without blocking antibody.

### HCV neutralization assays

HCV pseudoparticles (HCVpp) were generated by co-transfection of 293T cells with pNL4-3.lucR^-^E^-^ and the corresponding expression plasmids encoding the E1E2 from selected isolates with tier 1-4 antibody resistance^44–47^ at a 4:1 ratio by polyethylenimine (PEI, Polysciences). HCVpp neutralization was carried out on Huh-7 cells with mAbs (25 μg/ml) as recently described. ^48^ Virus infectivity was detected with Bright-Glo luciferase assay system (Promega), and percent neutralization was calculated as the virus infectivity inhibited at the antibody concentrations divided by the infectivity without antibody after background subtraction. The background infectivity of the pseudoparticle virus was defined by infecting cells with virus made only with pNL4-3.lucR-E-. Pseudoparticles displaying the vesicular stomatitis virus envelope glycoprotein G (VSVpp) were used as control for nonspecific neutralizing activity.

### Epitope mapping by alanine scanning mutagenesis

High-throughput scanning mutagenesis using an HCV E1E2 alanine scanning library (genotype 1a, strain H77; reference sequence NC_004102) was performed as described previously.^30^ Binding of the mAb to an individual alanine mutant was expressed as a % of the mAb’s binding signal obtained with the wild type E1E2.

### Antigen protein expression and purification

The SpyE1E2, E2ecto or E2c3 proteins (Table S2) were expressed and purified as previously described.^26,28^ The purified proteins were stored in 20 mM Tris pH 8.0 and 150 mM NaCl. The purity and integrity of recombinant proteins were confirmed by reducing and nonreducing SDS-PAGE.

### Biolayer interferometry

E2ecto, E2c3 or SpyE1E2 glycoprotein binding to antibodies was evaluated by BLI using an Octet RED96 instrument (ForteBio) as previously described.^8^ To compare the binding of antibodies to E2ecto and SpyE1E2 proteins, 10 μg/mL of antibodies are immobilized on Fab-2G biosensors. The bound antibody sensors were then dipped into the well with 20 μg of E2ecto or SpyE1E2 antigen proteins for 300s and moved to the well with 1× kinetics buffer to allow the dissociation of antigens from antibodies for 300s.

For the kinetic assay, antibodies at 10 μg/mL in 1× kinetics buffer (1× PBS, pH 7.4, 0.01% BSA, and 0.002% Tween 20) were loaded onto Fab-2G biosensors and analyzed for binding to E2ecto or spyE1E2 antigen at serial concentrations (640 nM, 320 nM, 160 nM, 80 nM, 40 nM, and 0 nM). The kinetic assay consisted of the following steps: 1) baseline with buffer; 2) loading with antibody proteins; 3) wash of unbound antibodies with buffer; 4) baseline with buffer; 5) association with antigen proteins; and 6) dissociation with buffer. For estimating the dissociation constant (K_d_), a 1:1 binding model was used. All binding data were collected at 10 °C.

### Negative-stain electron microscopy (nsEM) and image processing

The nsEM analysis was performed by the Core Microscopy Facility at The Scripps Research Institute following the protocol as previously described.^26^ Briefly, two micrograms of HCV-1 SpyE1E2 was mixed with 2 μg of Fab in 200 μl PBS and incubated at 4°C overnight for forming complex. The complex samples were diluted to concentrations of 0.005 and 0.008 mg/ml and 8 μL of complex sample was blotted onto carbon-coated copper grids (400 mesh) with glow-discharge for 2 minutes. Next, excess sample was removed, and the grids were negatively stained with 2% uranyl formate for 2 min. Excess stain was wicked away, and the grids were allowed to dry. The complex samples were analyzed at 120 kV with a Talos L120C transmission electron microscope (TEM, Thermo Fisher), and images were acquired using a CETA 16 M CMOS camera under 73,000× magnification at a resolution of 1.93 Å/pixel and defocus of 0.5-2 µm. Computational analysis of the images was performed using the high-performance computing core facility at The Scripps Research Institute. Briefly, the nsEM images were converted to MRC format by EMAN2^49^ for further processing by cryoSPARC v4.3.0.^50^ Micrographs were contrasting transfer function (CTF)-corrected by patch CTF estimation. Particles were selected using a Blob/template picker and later extracted with a box size of 140 pixel for 2D classification, with 100 and 250 Å used as the minimum and maximum particle sizes in blob picking. For all the dimer/nAb complexes, 3D models were generated by *Ab initio* reconstruction and optimized by heterogeneous and homogeneous refinement. All nsEM and fitted structure images were generated using Chimera X.^51^ and footprint analyzing by PyMOL (The PyMOL Molecular Graphics System, Version 3.0 Schrödinger, LLC).

### Crystal structure determination

Hk6a E2c3 was mixed with K400 Fab and H77 E2c3 protein was mixed with RM3-26 Fab in a 1:1.2 molar ratio and incubated overnight at 4°C. The complex mixture was further purified by gel filtration (Superdex 200 column) to remove unbound Fab. All Fabs and Fab-E2c3 complexes were screened for crystallization using the sitting drop vapor diffusion method. Initial crystallization conditions were obtained from robotic crystallization trials using our Rigaku CrystalMation system. The RM3-26 Fab - H77 E2c3 complex crystal used for data collection was grown with a well solution of 0.2 M potassium thiocyanate and 20% (w/v) polyethylene glycol 3,350. The crystal for unliganded K595 Fab was grown from 0.2 M potassium sulfate, 20% (w/v) polyethylene glycol 3350. The crystal obtained from K400 Fab was grown in 0.08 M sodium acetate (pH 4.6), 0.16 M ammonium sulfate, 20% (v/v) glycerol, 20% (w/v) polyethylene glycol 4000.

The crystals were flash-cooled in 80% (v/v) well solution and 20% (v/v) ethylene glycol as cryoprotectant before rapid plunging into liquid nitrogen. Diffraction data were collected at APS beamline 23-ID-D, SSRL 12-1 , and NSLS-II 17-ID-2, and then integrated and scaled with HKL-2000 ^52^ or XDS^53^. Statistics of the data collection are summarized in Table S3. The structures were determined by molecular replacement (MR) using the program Phaser-MR in Phenix ^54^. The MR models for Fabs were modeled with Repertoire Builder (https://sysimm.ifrec.osaka-u.ac.jp/rep_builder/).^55^ The model template for H77 E2c3 was from 1a53 E2ecto (PDB: 6MEJ).^56^ All models were examined and modified with the program Coot^57^ to assess model fit and any potential clashes.^57^ Refinement was carried out with Phenix.^54^ The H77 E2c3 coordinates in complex with RM3-26 are incomplete due to weak density for some regions; the final model includes residues Pro490-Val515, Phe550-Ala566, Gly598-Arg630, and Arg639-Asn645. While the SEC-purified K400-E2c3 complex was set up for crystallization, crystals were obtained only for the isolated K400 Fab. Final refinement statistics for all structures are summarized in Table S3. The quality of the structures were analyzed using MolProbity.^58^

### Structural analysis

Epitope and paratope residues, their detailed interactions as well as the buried surface area on the E2 upon binding of Fab, were identified and calculated with the Protein Interfaces, Surfaces and Assemblies (PISA) server at the European Bioinformatics Institute (www.ebi.ac.uk/pdbe/pisa/).^59^ Structure figures were generated by MacPyMol (DeLano Scientific LLC). Fabs are renumbered according to the Kabat nomenclature.^60^

### Bioinformatics

Antibody sequences were submitted to IgBLAST (https://www.ncbi.nlm.nih.gov/igblast/) and ImMunoGeneTics information system (IMGT, http://www.imgt.org/) for gene identification and genetic assignment. Multiple-sequence alignments were performed using MegAlign Pro program in Lasergene 15.3 (DNAstar). Maximum-likelihood phylogenetic tree was constructed using MEGA X software.

## Data availability

The X-ray coordinates and structure factors have been deposited in the Research Collaboratory for Structural Bioinformatics (RCSB) Protein Data Bank under accession codes 9MSK (RM3-26–H77 E2c3 complex), 9YJS (K400 Fab), and 9YK6 (K595 Fab).

## Acknowledgments

GM/CA@APS has been funded by the National Cancer Institute (ACB-12002) and the National Institute of General Medical Sciences (AGM-12006, P30GM138396). This research used resources of the Advanced Photon Source; a U.S. Department of Energy (DOE) Office of Science User Facility operated for the DOE Office of Science by Argonne National Laboratory under Contract No. DE-AC02-06CH11357. Use of the Stanford Synchrotron Radiation Lightsource, SLAC National Accelerator Laboratory, is supported by the U.S. Department of Energy, Office of Science, Office of Basic Energy Sciences under Contract No. DE-AC02-76SF00515. The SSRL Structural Molecular Biology Program is supported by the DOE Office of Biological and Environmental Research, and by the National Institutes of Health, National Institute of General Medical Sciences (P30GM133894). The contents of this publication are solely the responsibility of the authors and do not necessarily represent the official views of NIGMS or NIH. This research used resources [insert beamline(s)] of the National Synchrotron Light Source II, a U.S. Department of Energy (DOE) Office of Science User Facility operated for the DOE Office of Science by Brookhaven National Laboratory under Contract No. DE-SC0012704. We acknowledge the support of NIH grants AI168251 (M.L., J.Z., I.A.W.), AI158193, AI168917 (M.L.), AI086230 (G.M.L.), and AI168048 (T.R.F.). The funding bodies had no role in the design or conclusions of this study.

## AUTHOR CONTRIBUTIONS

F.C. and M.L. designed and conceived the study; F.C, E.G., S.C.L. Y.K. and S.H. performed mAb characterization; Y.L. and L.H performed nsEM; Y.T.K.N, R.L.S., and L.U. performed structural analysis; T.R.F. and G.M.L. provided research reagents before publication; M.L., J.Z, and I.A.W provided funding support; F.C., Y.T.K.N and Y.L. drafted the figures and manuscript. All authors read, edited, and approved the final manuscript.

---

**Figure S1.**
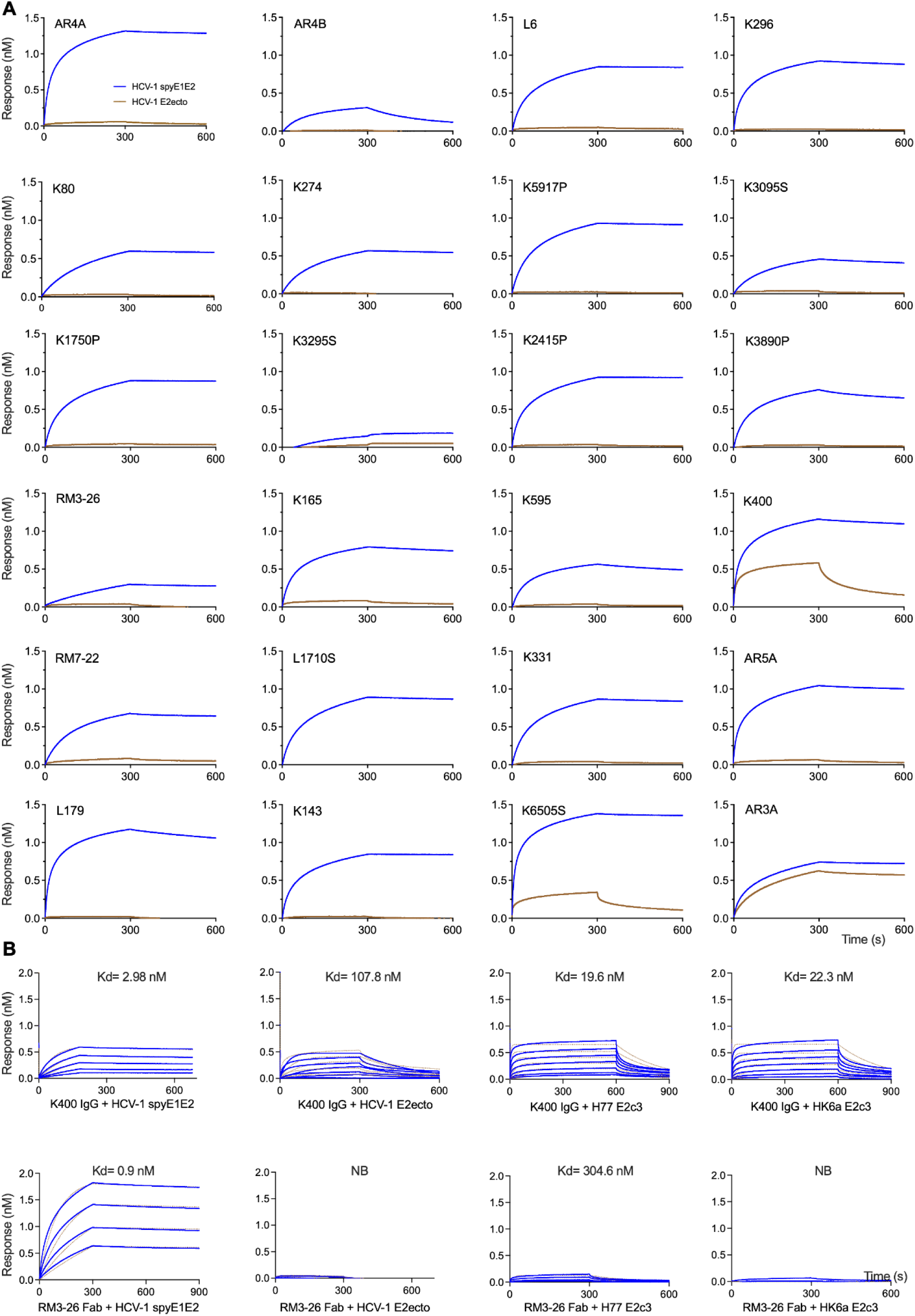
Biolayer interferometry (BLI) of representative BD antibodies to SpyE1E2 versus E2. (A). The antibodies were immobilized on Fab-2G biosensors. The HCV-1 E2ecto (brown) and SpyE1E2 (blue) proteins were used as analytes. The BLI binding response is shown in nanometers. (B) The binding kinetic curve and apparent Kd value of antibodies with E2 or E1E2 from different constructs. The response cure and fitting curves are shown in blue and brown, respectively.

**Figure S2.**
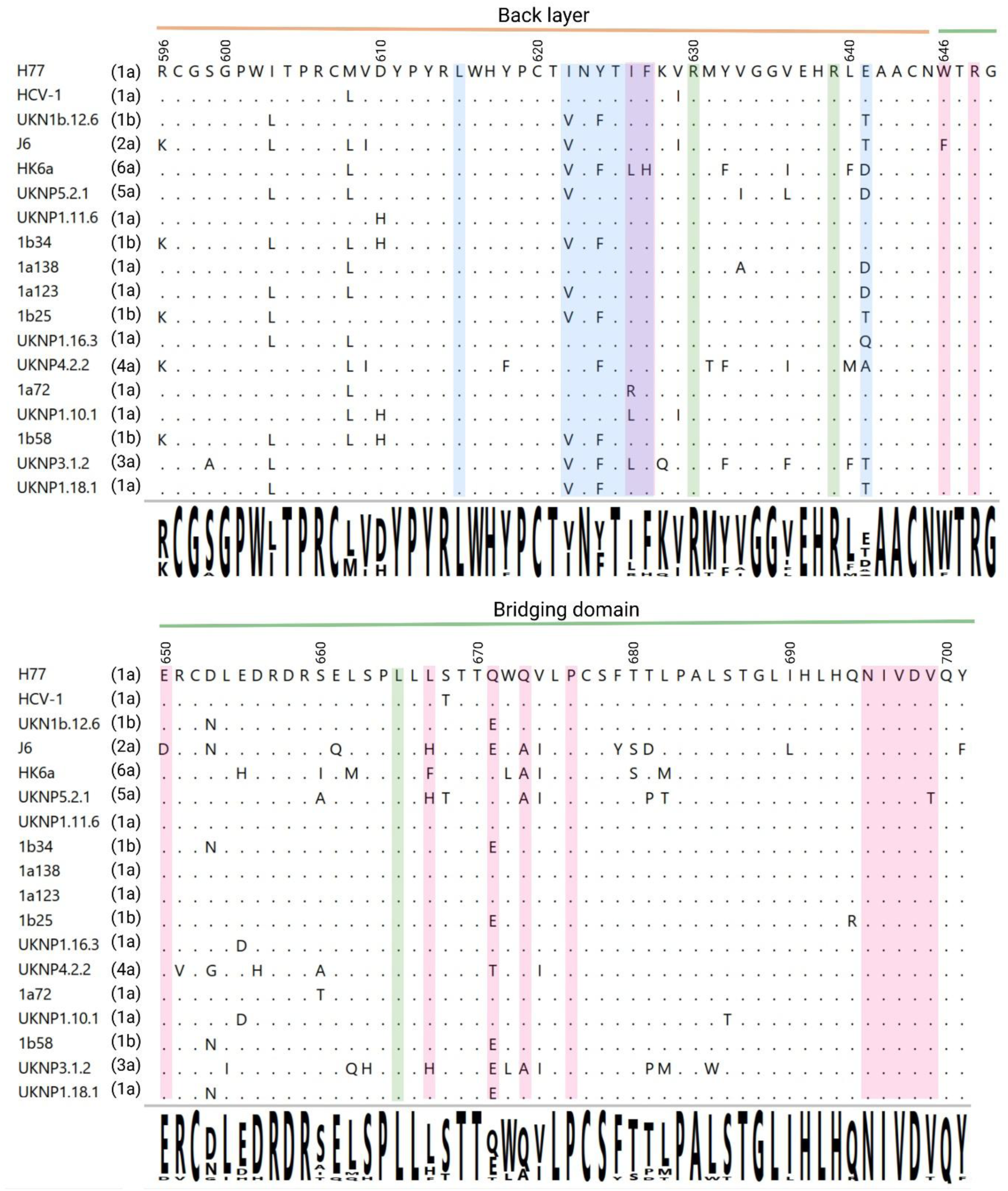
Sequence alignment of the back layer and BD regions from the HCV strains used in this study, related to. Figure 1C. Residues important for recognition by RM3-26, AR4A and AR5-specific antibodies are highlighted in blue, pink and green, respectively.

**Figure S3.**
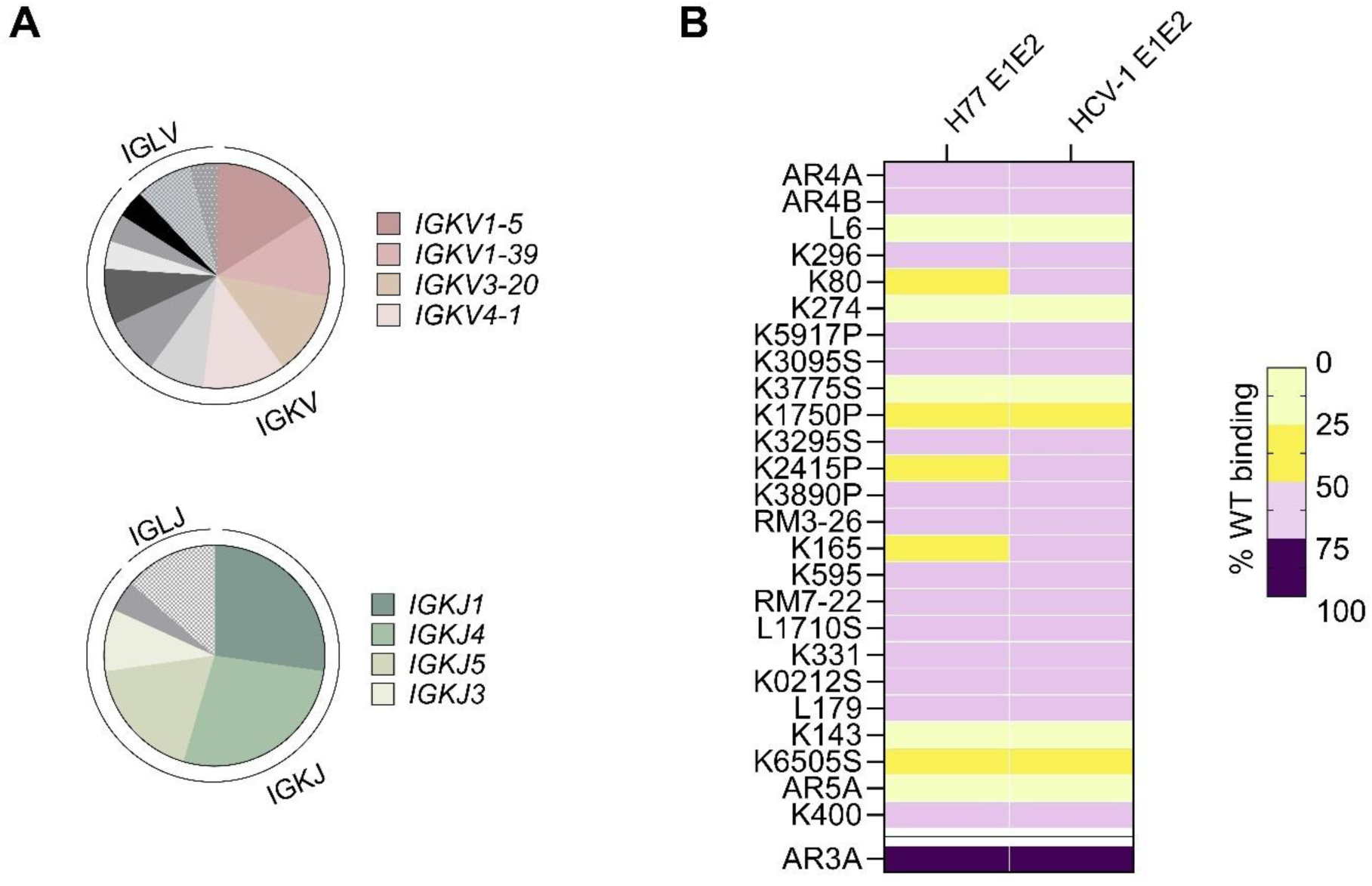
Light chain features of BD-targeting bnAbs. (A) Distribution of light chain variable (V) and joining (J) gene usage among BD-targeting bnAbs. (B) Binding activity of BD-targeting bnAbs in which wild-type heavy chains were paired with an unrelated light chain derived from a random influenza antibody.

**Figure S4.**
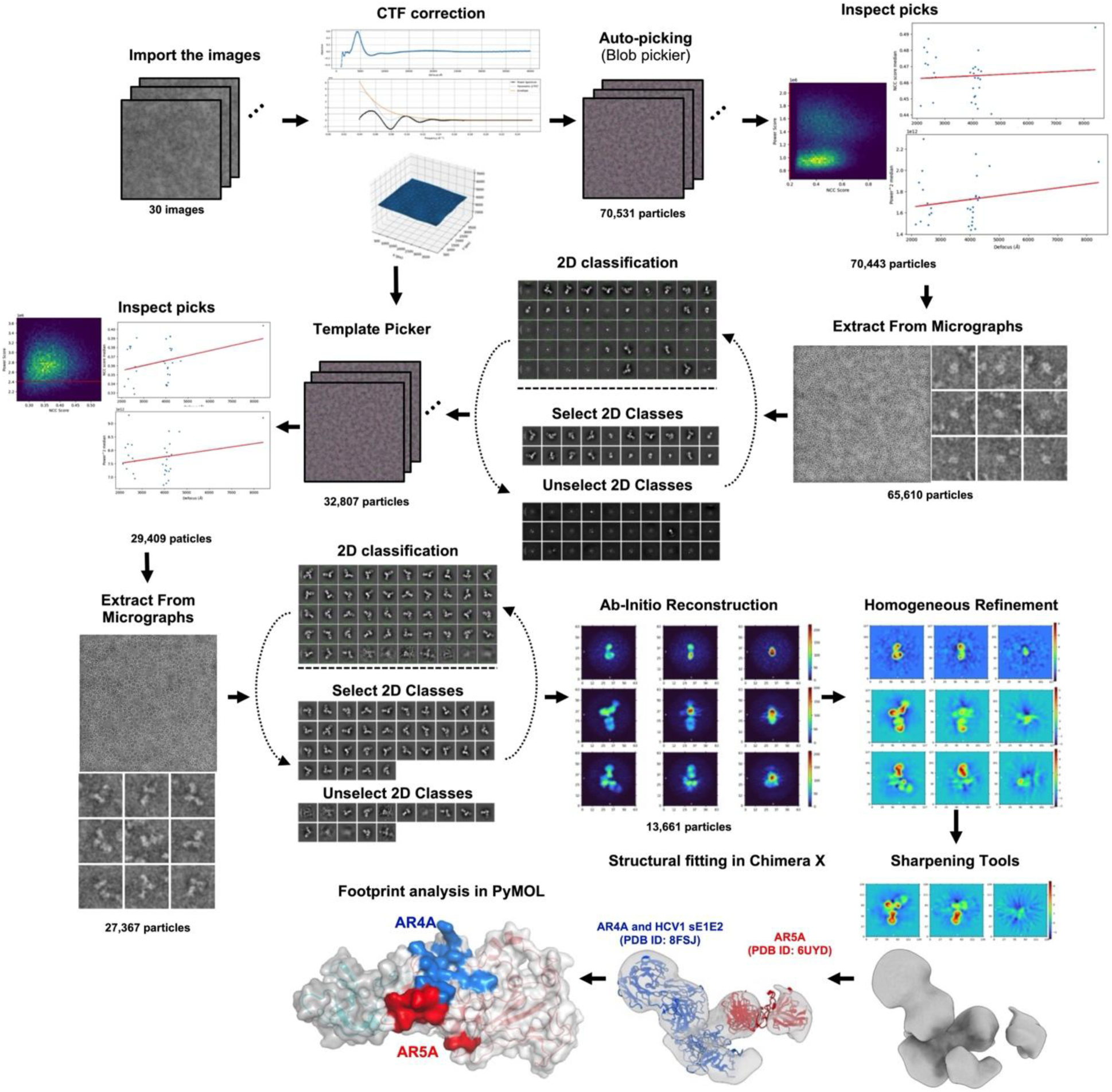
Schematic representation of image processing, 2D classification, 3D reconstruction of nsEM data, model fitting and epitope analysis obtained for SpyE1E2 complex in Fab using CryoSPARC. HCV-1 SpyE1E2 complexed with AR4A and AR5A Fabs is used as an example here.

**Figure S5.**
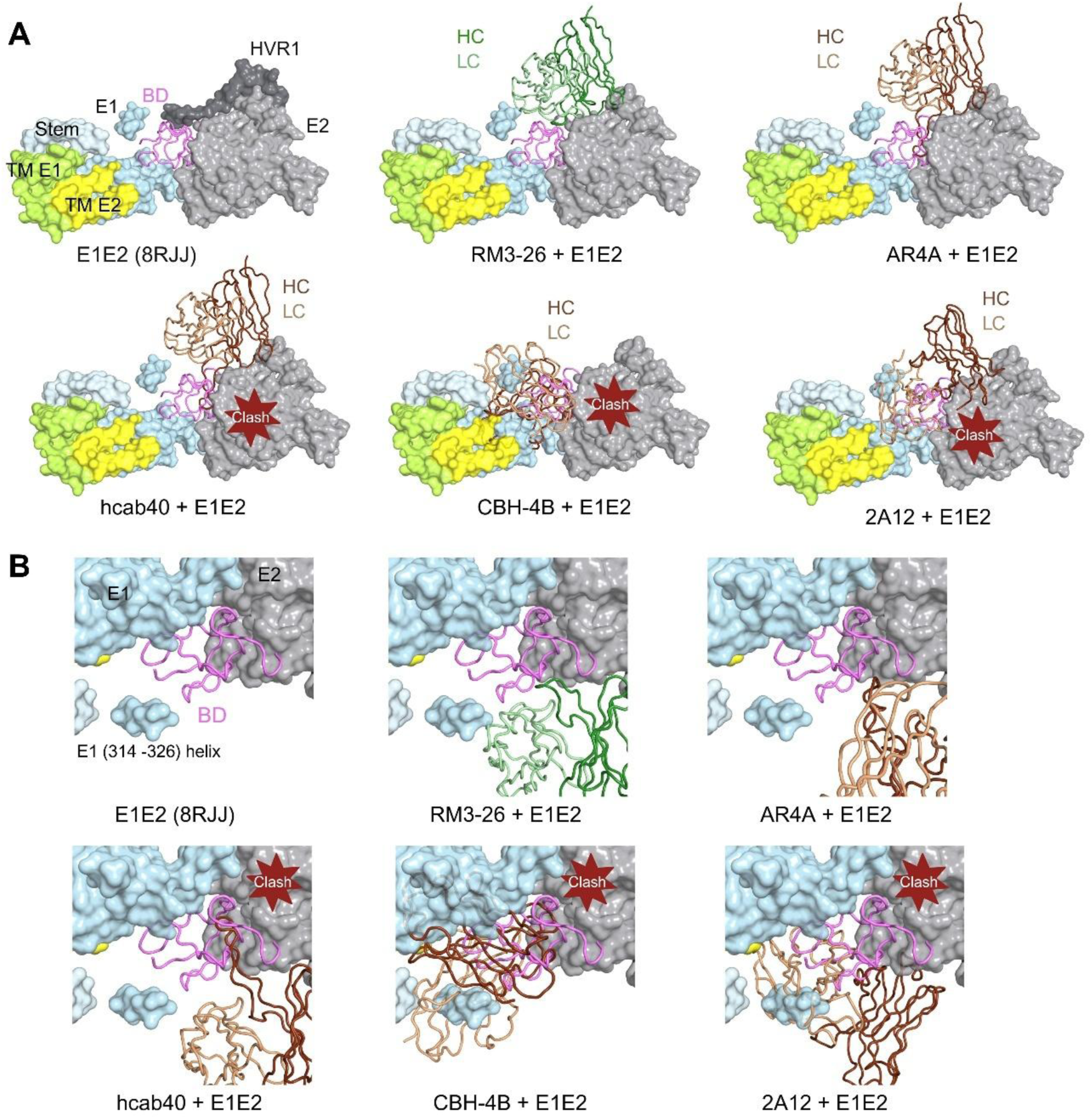
Modeling the interaction of back layer- and BD-targeting Abs on full-length E1E2. (A) Predicted model of RM3-26, hcab40, CBH4B, and AR4A bound to full-length E1E2 (PDB: 8RJJ). Each antibody-E2 complex was superimposed onto the full-length E1E2 (PDB: 8RJJ) based on the E2 region to generate the full-length complex model. The E2 HVR1 is shown in red, E2 BD in violet, E2 transmembrane (TM) in yellow, other E2 in vanadium. The E1 stem is shown in sky blue, E1 TM in lime green, and other E1 in cyan. The heavy and light chains (HC and LC) of RM3-26 are shown in dark and light green, respectively, while the heavy and light chains of other antibodies are shown in dark and light brown, respectively. (B) The zoomed-in view of Abs in complex with E1E2 model. Antibodies hcab40, CBH4B, and 2A12 appear to clash with E2 BD while RM3-26 and AR4A do not.

**Figure S6.**
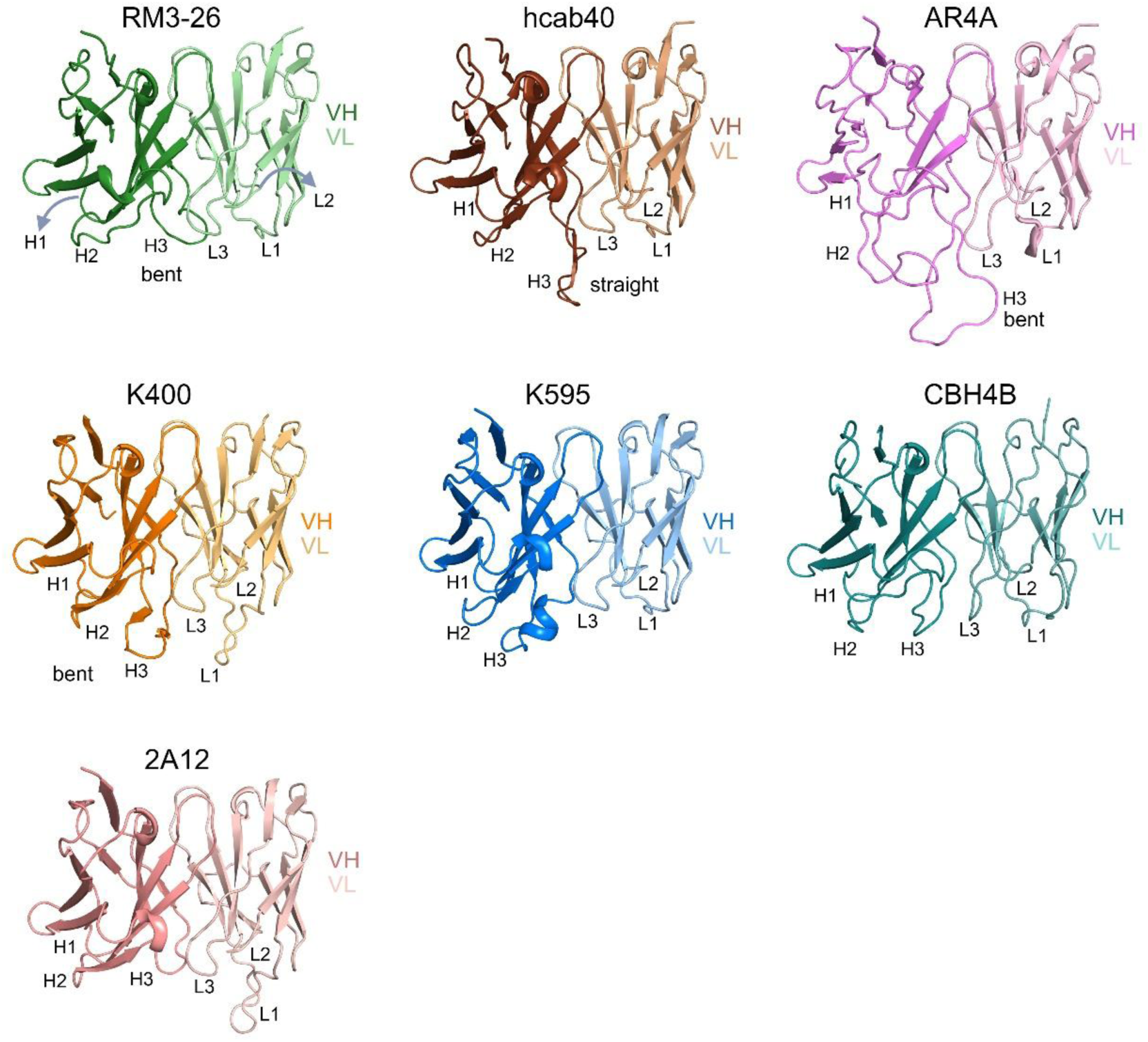
Structure of Fabs. The heavy (VL) and light (VL) chain variable domains of RM3-26, hcab40, AR4A, K400, K595, CBH4B, and 2A12 are shown as cartoons. Complementarity-determining regions (CDRs) H1-H3 and L1-L3 are indicated.

**Table S1.**
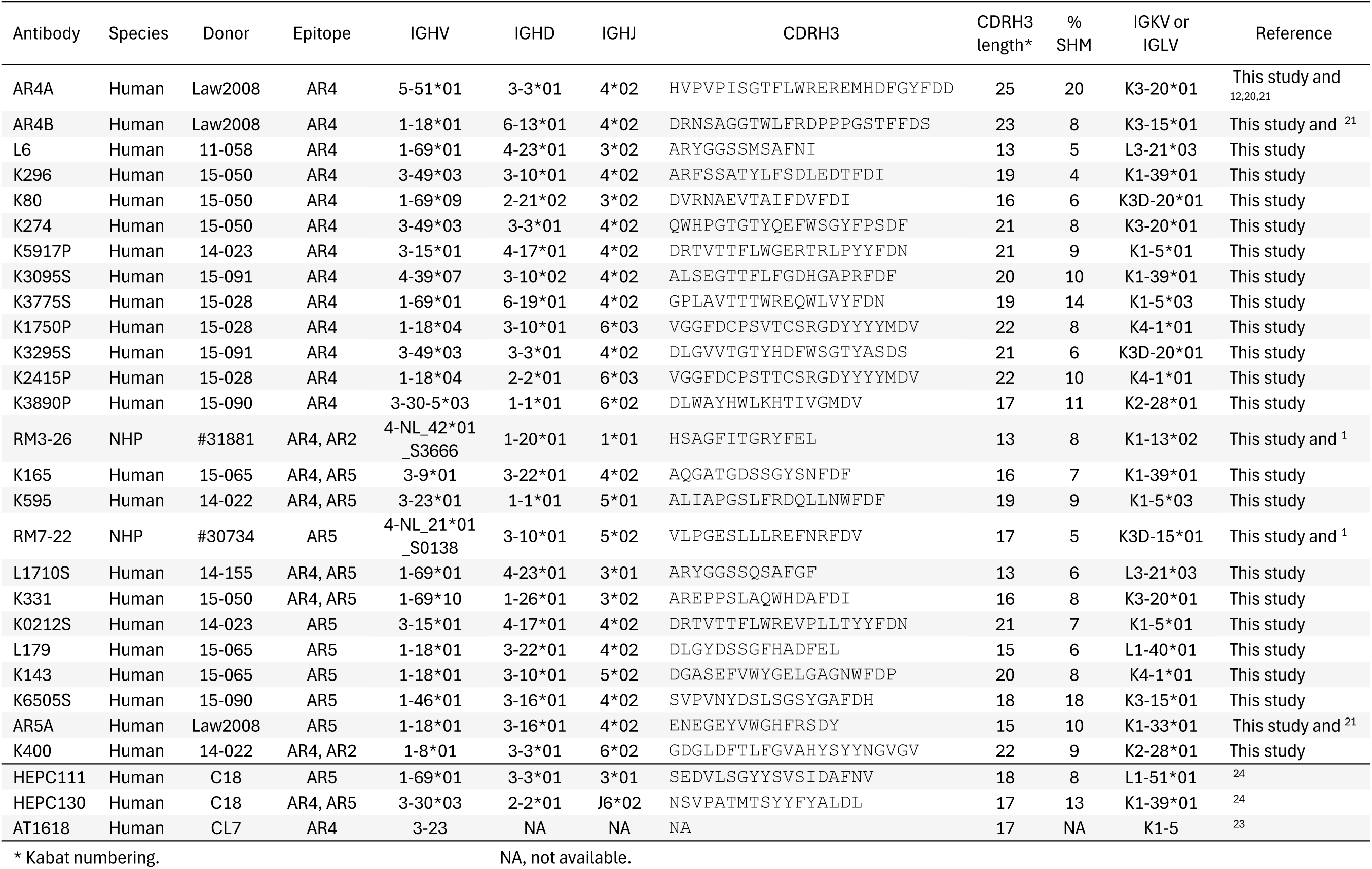
Sequence features of BD-targeting nAbs.

**Table S2.**
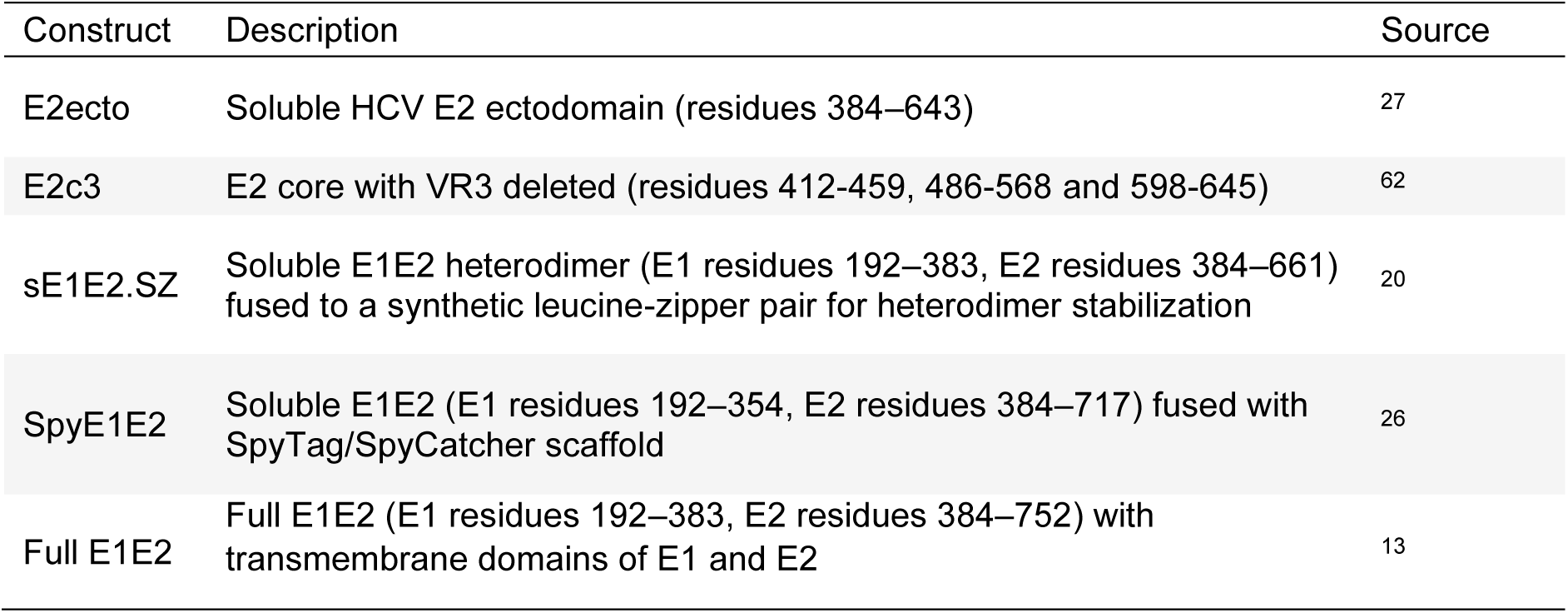
HCV protein constructs used in this study.

**Table S3.**
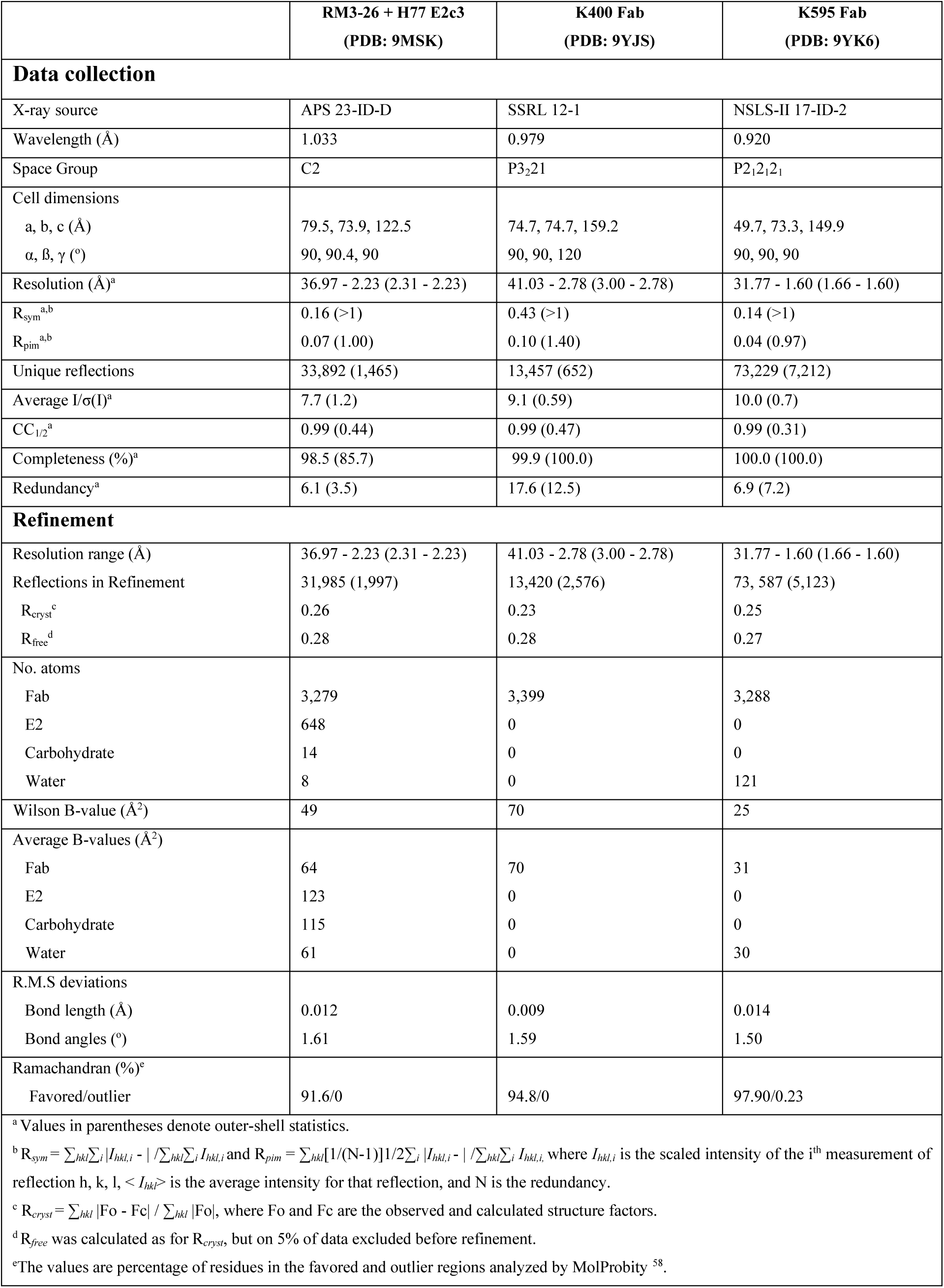
Data collection and refinement statistics of BD nAb crystal structures.

**Table S4.**
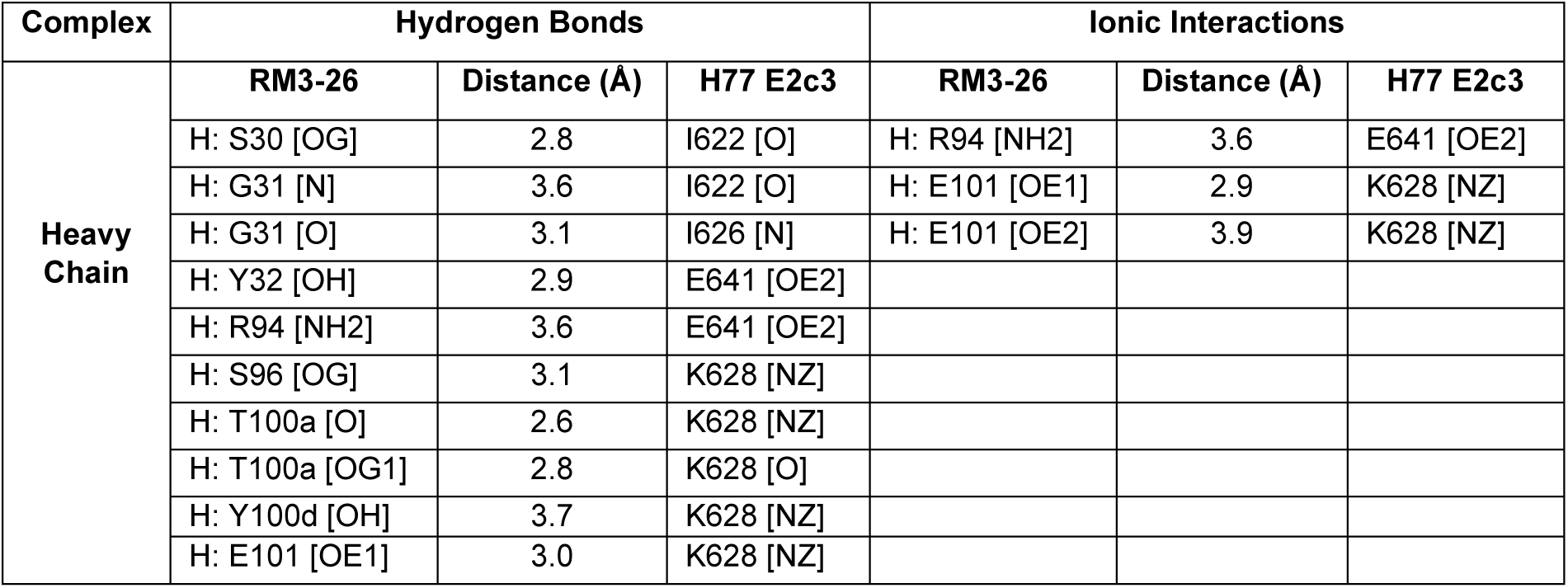
Polar and electrostatic interactions between Fab RM3-26 and H77 E2c3.

**Table S5.**
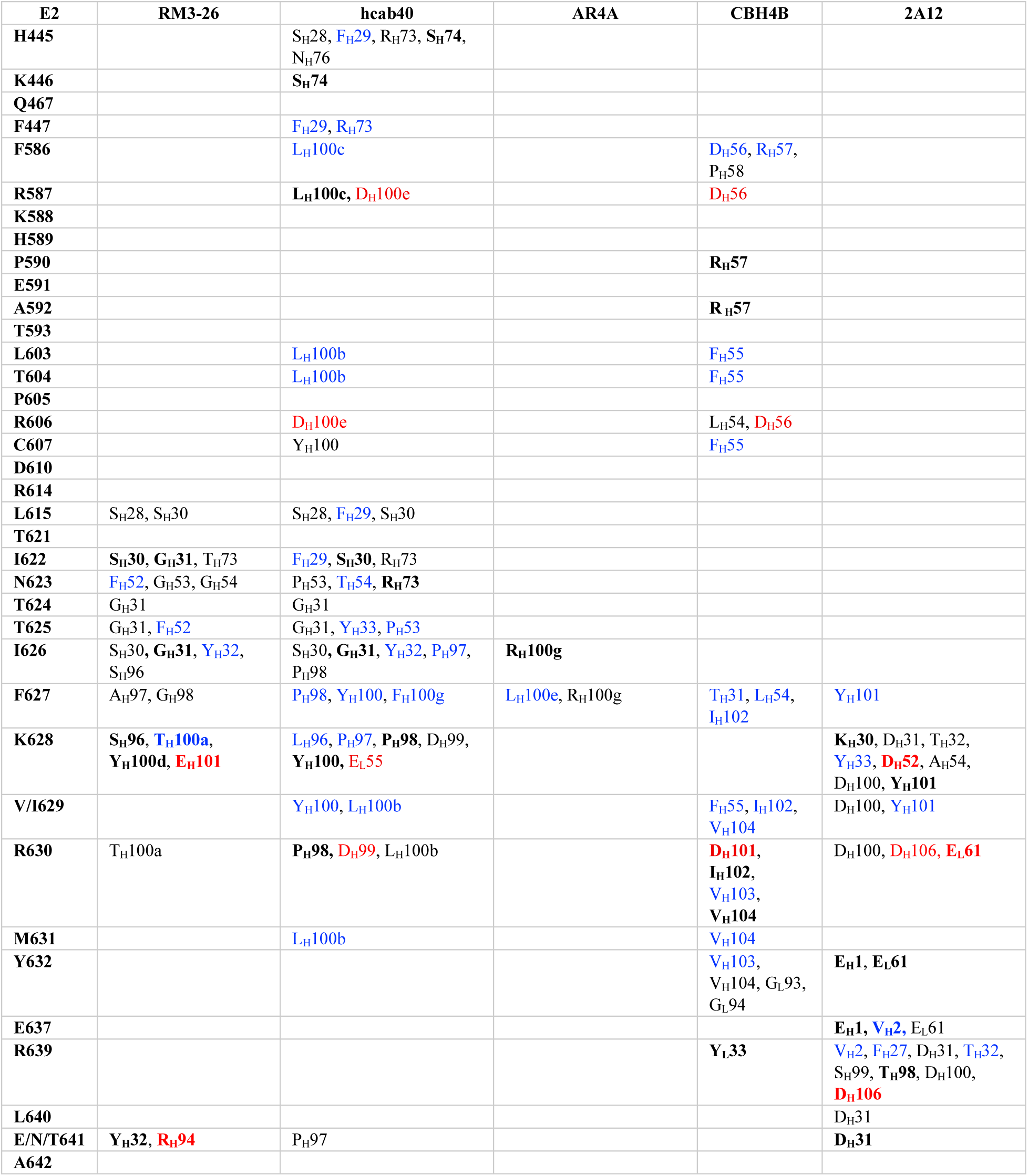
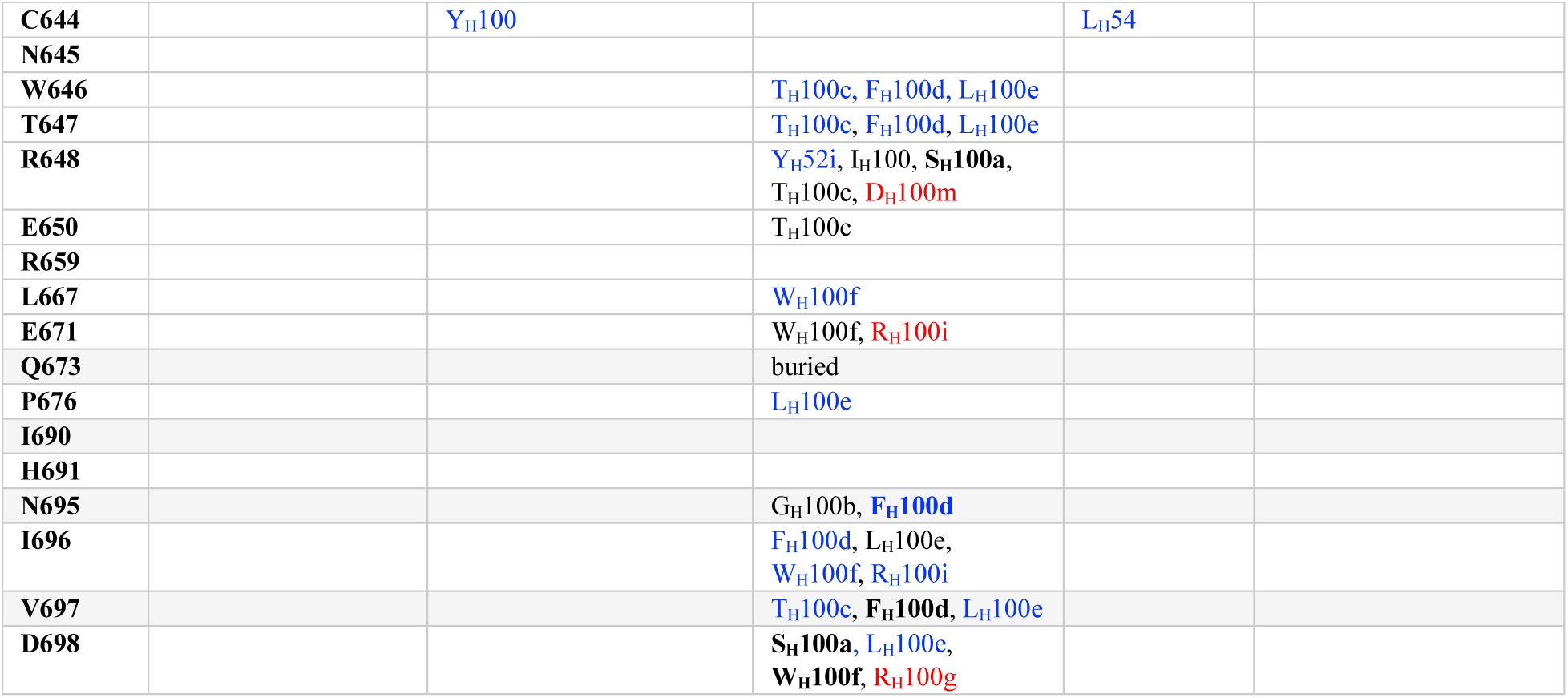
Back layer- and BD-targeting antibody interaction with E2. Red (ionic interaction), bold (hydrogen bonds), and blue (hydrophobic interactions). hcab40 in complex with 1b09 E2ecto (PDB 8WOX), AR4A in complex with 1b09 sE1E2. SZ (PDB 8FSJ), CBH4B in complex with 1b09 E2ecto (PDB: 8TXQ), and 2A12 in complex with J6 N-terminal truncated E2 (aa 456-713) (PDB: 8DK6) are used for the analysis.

## References

1. Chen, F., Nagy, K., Chavez, D., Willis, S., McBride, R., Giang, E., Honda, A., Bukh, J., Ordoukhanian, P., Zhu, J., et al. (2020). Antibody responses to immunization with HCV envelope glycoproteins as a baseline for B-cell-based vaccine development. Gastroenterology 158, 1058–1071 e1056. 10.1053/j.gastro.2019.11.282.

2. Page, K., Melia, M.T., Veenhuis, R.T., Winter, M., Rousseau, K.E., Massaccesi, G., Osburn, W.O., Forman, M., Thomas, E., Thornton, K., et al. (2021). Randomized Trial of a Vaccine Regimen to Prevent Chronic HCV Infection. N. Engl. J. Med. 384, 541–549. 10.1056/NEJMoa2023345.

3. Smith, D.B., Bukh, J., Kuiken, C., Muerhoff, A.S., Rice, C.M., Stapleton, J.T., and Simmonds, P. (2014). Expanded classification of hepatitis C virus into 7 genotypes and 67 subtypes: updated criteria and genotype assignment web resource. Hepatology 59, 318–327. 10.1002/hep.26744.

4. Lavie, M., Hanoulle, X., and Dubuisson, J. (2018). Glycan Shielding and Modulation of Hepatitis C Virus Neutralizing Antibodies. Front. Immunol. 9, 910. 10.3389/fimmu.2018.00910.

5. Sepulveda-Crespo, D., Resino, S., and Martinez, I. (2020). Hepatitis C virus vaccine design: focus on the humoral immune response. J. Biomed. Sci. 27, 78. 10.1186/s12929-020-00669-4.

6. de Jong, Y.P., Dorner, M., Mommersteeg, M.C., Xiao, J.W., Balazs, A.B., Robbins, J.B., Winer, B.Y., Gerges, S., Vega, K., Labitt, R.N., et al. (2014). Broadly neutralizing antibodies abrogate established hepatitis C virus infection. Sci. Transl. Med. 6, 254ra129. 10.1126/scitranslmed.3009512.

7. Osburn, W.O., Snider, A.E., Wells, B.L., Latanich, R., Bailey, J.R., Thomas, D.L., Cox, A.L., and Ray, S.C. (2014). Clearance of hepatitis C infection is associated with the early appearance of broad neutralizing antibody responses. Hepatology 59, 2140–2151. 10.1002/hep.27013.

8. Chen, F., Tzarum, N., Lin, X., Giang, E., Velazquez-Moctezuma, R., Augestad, E.H., Nagy, K., He, L., Hernandez, M., Fouch, M.E., et al. (2021). Functional convergence of a germline-encoded neutralizing antibody response in rhesus macaques immunized with HCV envelope glycoproteins. Immunity 54, 781–796 e784. 10.1016/j.immuni.2021.02.013.

9. Logvinoff, C., Major, M.E., Oldach, D., Heyward, S., Talal, A., Balfe, P., Feinstone, S.M., Alter, H., Rice, C.M., and McKeating, J.A. (2004). Neutralizing antibody response during acute and chronic hepatitis C virus infection. Proc. Natl. Acad. Sci. U. S. A. 101, 10149–10154. 10.1073/pnas.0403519101.

10. Merat, S.J., Molenkamp, R., Wagner, K., Koekkoek, S.M., van de Berg, D., Yasuda, E., Bohne, M., Claassen, Y.B., Grady, B.P., Prins, M., et al. (2016). Hepatitis C virus broadly neutralizing monoclonal antibodies isolated 25 years after spontaneous clearance. PLoS One 11, e0165047. 10.1371/journal.pone.0165047.

11. Bailey, J.R., Flyak, A.I., Cohen, V.J., Li, H., Wasilewski, L.N., Snider, A.E., Wang, S., Learn, G.H., Kose, N., Loerinc, L., et al. (2017). Broadly neutralizing antibodies with few somatic mutations and hepatitis c virus clearance. JCI Insight 2, e92872. 10.1172/jci.insight.92872.

12. Torrents de la Pena, A., Sliepen, K., Eshun-Wilson, L., Newby, M.L., Allen, J.D., Zon, I., Koekkoek, S., Chumbe, A., Crispin, M., Schinkel, J., et al. (2022). Structure of the hepatitis C virus E1E2 glycoprotein complex. Science 378, 263–269. 10.1126/science.abn9884.

13. Augestad, E.H., Holmboe Olesen, C., Gronberg, C., Soerensen, A., Velazquez-Moctezuma, R., Fanalista, M., Bukh, J., Wang, K., Gourdon, P., and Prentoe, J. (2024). The hepatitis C virus envelope protein complex is a dimer of heterodimers. Nature. 10.1038/s41586-024-07783-5.

14. Lindenbach, B.D., and Rice, C.M. (2013). The ins and outs of hepatitis C virus entry and assembly. Nat. Rev. Microbiol. 11, 688–700. 10.1038/nrmicro3098.

15. Tong, Y., Lavillette, D., Li, Q., and Zhong, J. (2018). Role of Hepatitis C Virus Envelope Glycoprotein E1 in Virus Entry and Assembly. Front. Immunol. 9, 1411. 10.3389/fimmu.2018.01411.

16. Tzarum, N., Wilson, I.A., and Law, M. (2018). The neutralizing face of hepatitis C virus E2 envelope glycoprotein. Front. Immunol. 9, 1315. 10.3389/fimmu.2018.01315.

17. Law, M., Maruyama, T., Lewis, J., Giang, E., Tarr, A.W., Stamataki, Z., Gastaminza, P., Chisari, F.V., Jones, I.M., Fox, R.I., et al. (2008). Broadly neutralizing antibodies protect against hepatitis c virus quasispecies challenge. Nat. Med. 14, 25–27. 10.1038/nm1698.

18. Weber, T., Potthoff, J., Bizu, S., Labuhn, M., Dold, L., Schoofs, T., Horning, M., Ercanoglu, M.S., Kreer, C., Gieselmann, L., et al. (2022). Analysis of antibodies from HCV elite neutralizers identifies genetic determinants of broad neutralization. Immunity 55, 341–354 e347. 10.1016/j.immuni.2021.12.003.

19. Chen, F., Tzarum, N., Wilson, I.A., and Law, M. (2019). V_H_1-69 antiviral broadly neutralizing antibodies: Genetics, structures, and relevance to rational vaccine design. Curr. Opin. Virol. 34, 149–159. 10.1016/j.coviro.2019.02.004.

20. Metcalf, M.C., Janus, B.M., Yin, R., Wang, R., Guest, J.D., Pozharski, E., Law, M., Mariuzza, R.A., Toth, E.A., Pierce, B.G., et al. (2023). Structure of engineered hepatitis C virus E1E2 ectodomain in complex with neutralizing antibodies. Nat Commun 14, 3980. 10.1038/s41467-023-39659-z.

21. Giang, E., Dorner, M., Prentoe, J.C., Dreux, M., Evans, M.J., Bukh, J., Rice, C.M., Ploss, A., Burton, D.R., and Law, M. (2012). Human broadly neutralizing antibodies to the envelope glycoprotein complex of hepatitis C virus. Proc. Natl. Acad. Sci. U. S. A. 109, 6205–6210. 10.1073/pnas.1114927109.

22. Chen, F., Giang, E., Natarajan, P., Mondala, T.S., Head, S.R., Lau, S.C., Sundaresan, A., Kulakova, L., Lin, X., Aneja, J., et al. (2025). B cell transcriptomics reveals lasting dysregulation and rapid decline of protective immune memory after chronic hepatitis C cure. bioRxiv, 2025.2010.2024.664545. 10.1101/2025.10.24.664545.

23. Merat, S.J., Bru, C., van de Berg, D., Molenkamp, R., Tarr, A.W., Koekkoek, S., Kootstra, N.A., Prins, M., Ball, J.K., Bakker, A.Q., et al. (2019). Cross-genotype AR3-specific neutralizing antibodies confer long-term protection in injecting drug users after hcv clearance. J. Hepatol. 71, 14–24. 10.1016/j.jhep.2019.02.013.

24. Colbert, M.D., Flyak, A.I., Ogega, C.O., Kinchen, V.J., Massaccesi, G., Hernandez, M., Davidson, E., Doranz, B.J., Cox, A.L., Crowe, J.E., Jr., and Bailey, J.R. (2019). Broadly neutralizing antibodies targeting new sites of vulnerability in hepatitis c virus E1E2. J. Virol. 93. 10.1128/JVI.02070-18.

25. Guest, J.D., Wang, R., Elkholy, K.H., Chagas, A., Chao, K.L., Cleveland, T.E.t., Kim, Y.C., Keck, Z.Y., Marin, A., Yunus, A.S., et al. (2021). Design of a native-like secreted form of the hepatitis C virus E1E2 heterodimer. Proc. Natl. Acad. Sci. U. S. A. 118. 10.1073/pnas.2015149118.

26. He, L., Lee, Y.Z., Zhang, Y.N., Newby, M.L., Janus, B.M., Gonzalez, F.G., Ward, G., DesRoberts, C., Hung, S.H., Giang, E., et al. (2025). Native-like soluble E1E2 glycoprotein heterodimers on self-assembling protein nanoparticles for hepatitis C virus vaccine design. bioRxiv. 10.1101/2025.05.16.654559.

27. Khan, A.G., Whidby, J., Miller, M.T., Scarborough, H., Zatorski, A.V., Cygan, A., Price, A.A., Yost, S.A., Bohannon, C.D., Jacob, J., et al. (2014). Structure of the core ectodomain of the hepatitis C virus envelope glycoprotein 2. Nature 509, 381–384. 10.1038/nature13117.

28. Tzarum, N., Giang, E., Kadam, R.U., Chen, F., Nagy, K., Augestad, E.H., Velazquez-Moctezuma, R., Keck, Z.Y., Hua, Y., Stanfield, R.L., et al. (2020). An alternate conformation of HCV E2 neutralizing face as an additional vaccine target. Sci Adv 6, eabb5642. 10.1126/sciadv.abb5642.

29. Dubuisson, J., Hsu, H.H., Cheung, R.C., Greenberg, H.B., Russell, D.G., and Rice, C.M. (1994). Formation and intracellular localization of hepatitis C virus envelope glycoprotein complexes expressed by recombinant vaccinia and Sindbis viruses. J. Virol. 68, 6147–6160. 10.1128/JVI.68.10.6147-6160.1994.

30. Gopal, R., Jackson, K., Tzarum, N., Kong, L., Ettenger, A., Guest, J., Pfaff, J.M., Barnes, T., Honda, A., Giang, E., et al. (2017). Probing the antigenicity of hepatitis c virus envelope glycoprotein complex by high-throughput mutagenesis. PLoS Pathog. 13, e1006735. 10.1371/journal.ppat.1006735.

31. Kong, L., Giang, E., Nieusma, T., Kadam, R.U., Cogburn, K.E., Hua, Y., Dai, X., Stanfield, R.L., Burton, D.R., Ward, A.B., et al. (2013). Hepatitis C virus E2 envelope glycoprotein core structure. Science 342, 1090–1094. 10.1126/science.1243876.

32. Tzarum, N., Giang, E., Kong, L., He, L., Prentoe, J., Augestad, E., Hua, Y., Castillo, S., Lauer, G.M., Bukh, J., et al. (2019). Genetic and structural insights into broad neutralization of hepatitis c virus by human V_H_1-69 antibodies. Sci Adv 5, eaav1882. 10.1126/sciadv.aav1882.

33. He, L., Tzarum, N., Lin, X., Shapero, B., Sou, C., Mann, C.J., Stano, A., Zhang, L., Nagy, K., Giang, E., et al. (2020). Proof of concept for rational design of Hepatitis C virus E2 core nanoparticle vaccines. Sci. Adv. 6, eaaz6225. 10.1126/sciadv.aaz6225.

34. Ogega, C.O., Skinner, N.E., Schoenle, M.V., Wilcox, X.E., Frumento, N., Wright, D.A., Paul, H.T., Sinnis-Bourozikas, A., Clark, K.E., Figueroa, A., et al. (2024). Convergent evolution and targeting of diverse E2 epitopes by human broadly neutralizing antibodies are associated with HCV clearance. Immunity 57, 890–903.e896. 10.1016/j.immuni.2024.03.001.

35. Kumar, A., Rohe, T.C., Elrod, E.J., Khan, A.G., Dearborn, A.D., Kissinger, R., Grakoui, A., and Marcotrigiano, J. (2023). Regions of Hepatitis C virus E2 required for membrane association. Nat. Commun. 14, 433. 10.1038/s41467-023-36183-y.

36. Shahid, S., Karade, S.S., Hasan, S.S., Yin, R., Jiang, L., Liu, Y., Felbinger, N., Kulakova, L., Toth, E.A., Keck, Z.Y., et al. (2025). Cryo-EM structures of HCV E2 glycoprotein bound to neutralizing and non-neutralizing antibodies determined using bivalent Fabs as fiducial markers. Commun Biol 8, 825. 10.1038/s42003-025-08239-w.

37. Salas, J.H., Urbanowicz, R.A., Guest, J.D., Frumento, N., Figueroa, A., Clark, K.E., Keck, Z., Cowton, V.M., Cole, S.J., Patel, A.H., et al. (2022). An Antigenically Diverse, Representative Panel of Envelope Glycoproteins for Hepatitis C Virus Vaccine Development. Gastroenterology 162, 562–574. 10.1053/j.gastro.2021.10.005.

38. Flyak, A.I., Ruiz, S.E., Salas, J., Rho, S., Bailey, J.R., and Bjorkman, P.J. (2020). An ultralong CDRH2 in HCV neutralizing antibody demonstrates structural plasticity of antibodies against E2 glycoprotein. Elife 9. 10.7554/eLife.53169.

39. Klein, F., Diskin, R., Scheid, J.F., Gaebler, C., Mouquet, H., Georgiev, I.S., Pancera, M., Zhou, T., Incesu, R.B., Fu, B.Z., et al. (2013). Somatic mutations of the immunoglobulin framework are generally required for broad and potent hiv-1 neutralization. Cell 153, 126–138. 10.1016/j.cell.2013.03.018.

40. Mankowski, M.C., Kinchen, V.J., Wasilewski, L.N., Flyak, A.I., Ray, S.C., Crowe, J.E., Jr., and Bailey, J.R. (2018). Synergistic anti-hcv broadly neutralizing human monoclonal antibodies with independent mechanisms. Proc. Natl. Acad. Sci. U. S. A. 115, E82–E91. 10.1073/pnas.1718441115.

41. Radic, L., Offersgaard, A., Kadava, T., Zon, I., Capella-Pujol, J., Mulder, F., Koekkoek, S., Spek, V., Chumbe, A., Bukh, J., et al. (2025). Bispecific antibodies against the hepatitis C virus E1E2 envelope glycoprotein. Proc. Natl. Acad. Sci. U. S. A. 122, e2420402122. 10.1073/pnas.2420402122.

42. Wang, R., Suzuki, S., Guest, J.D., Heller, B., Almeda, M., Andrianov, A.K., Marin, A., Mariuzza, R.A., Keck, Z.Y., Foung, S.K.H., et al. (2022). Induction of broadly neutralizing antibodies using a secreted form of the hepatitis C virus E1E2 heterodimer as a vaccine candidate. Proc. Natl. Acad. Sci. U. S. A. 119, e2112008119. 10.1073/pnas.2112008119.

43. Weaver, G.C., Villar, R.F., Kanekiyo, M., Nabel, G.J., Mascola, J.R., and Lingwood, D. (2016). In vitro reconstitution of B cell receptor-antigen interactions to evaluate potential vaccine candidates. Nat. Protoc. 11, 193–213. 10.1038/nprot.2016.009.

44. Bartosch, B., Dubuisson, J., and Cosset, F.L. (2003). Infectious hepatitis c virus pseudo-particles containing functional e1-e2 envelope protein complexes. J. Exp. Med. 197, 633–642.

45. Lavillette, D., Tarr, A.W., Voisset, C., Donot, P., Bartosch, B., Bain, C., Patel, A.H., Dubuisson, J., Ball, J.K., and Cosset, F.L. (2005). Characterization of host-range and cell entry properties of the major genotypes and subtypes of hepatitis c virus. Hepatology 41, 265–274. 10.1002/hep.20542.

46. Li, Y.-P., Ramirez, S., Mikkelsen, L., and Bukh, J. (2015). Efficient infectious cell culture systems of the hepatitis C virus (HCV) prototype strains HCV-1 and H77. Journal of Virology 89, 811–823. 10.1128/jvi.02877-14.

47. Meunier, J.-C., Engle, R.E., Faulk, K., Zhao, M., Bartosch, B., Alter, H., Emerson, S.U., Cosset, F.-L., Purcell, R.H., and Bukh, J. (2005). Evidence for cross-genotype neutralization of hepatitis C virus pseudo-particles and enhancement of infectivity by apolipoprotein C1. Proceedings of the National Academy of Sciences of the United States of America 102, 4560–4565. 10.1073/pnas.0501275102.

48. Bailey, J.R., Urbanowicz, R.A., Ball, J.K., Law, M., and Foung, S.K.H. (2019). Standardized method for the study of antibody neutralization of HCV pseudoparticles (HCVpp). Methods Mol. Biol. 1911, 441–450. 10.1007/978-1-4939-8976-8_30.

49. Tang, G., Peng, L., Baldwin, P.R., Mann, D.S., Jiang, W., Rees, I., and Ludtke, S.J. (2007). EMAN2: an extensible image processing suite for electron microscopy. J. Struct. Biol. 157, 38–46. 10.1016/j.jsb.2006.05.009.

50. Punjani, A., Rubinstein, J.L., Fleet, D.J., and Brubaker, M.A. (2017). cryoSPARC: algorithms for rapid unsupervised cryo-EM structure determination. Nat Methods 14, 290–296. 10.1038/nmeth.4169.

51. Meng, E.C., Goddard, T.D., Pettersen, E.F., Couch, G.S., Pearson, Z.J., Morris, J.H., and Ferrin, T.E. (2023). UCSF ChimeraX: Tools for structure building and analysis. Protein Sci. 32, e4792. 10.1002/pro.4792.

52. Otwinowski, Z., and Minor, W. (1997). Processing of X-ray diffraction data collected in oscillation mode. Methods Enzymol. 276, 307–326. 10.1016/s0076-6879(97)76066-x.

53. Kabsch, W. (2010). XDS. Acta Crystallogr D Biol Crystallogr 66, 125–132. 10.1107/s0907444909047337.

54. Adams, P.D., Grosse-Kunstleve, R.W., Hung, L.W., Ioerger, T.R., McCoy, A.J., Moriarty, N.W., Read, R.J., Sacchettini, J.C., Sauter, N.K., and Terwilliger, T.C. (2002). PHENIX: Building new software for automated crystallographic structure determination. Acta Crystallogr. D Biol. Crystallogr. 58, 1948–1954. 10.1107/s0907444902016657.

55. Schritt, D., Li, S., Rozewicki, J., Katoh, K., Yamashita, K., Volkmuth, W., Cavet, G., and Standley, D.M. (2019). Repertoire builder: High-throughput structural modeling of B and T cell receptors. Mol. Syst. Des. Eng. 4, 761–768. 10.1039/C9ME00020H.

56. Flyak, A.I., Ruiz, S., Colbert, M.D., Luong, T., Crowe, J.E., Jr., Bailey, J.R., and Bjorkman, P.J. (2018). HCV broadly neutralizing antibodies use a CDRH3 disulfide motif to recognize an E2 glycoprotein site that can be targeted for vaccine design. Cell Host & Microbe 24, 703–716.e703. 10.1016/j.chom.2018.10.009.

57. Emsley, P., and Cowtan, K. (2004). Coot: Model-building tools for molecular graphics. Acta Crystallogr. D Biol. Crystallogr. 60, 2126–2132. 10.1107/s0907444904019158.

58. Chen, V.B., Arendall, W.B., 3rd, Headd, J.J., Keedy, D.A., Immormino, R.M., Kapral, G.J., Murray, L.W., Richardson, J.S., and Richardson, D.C. (2010). MolProbity: All-atom structure validation for macromolecular crystallography. Acta Crystallogr. D Biol. Crystallogr. 66, 12–21. 10.1107/s0907444909042073.

59. Krissinel, E., and Henrick, K. (2007). Inference of macromolecular assemblies from crystalline state. J Mol Biol 372, 774–797. 10.1016/j.jmb.2007.05.022.

60. Abhinandan, K.R., and Martin, A.C. (2008). Analysis and improvements to Kabat and structurally correct numbering of antibody variable domains. Mol Immunol 45, 3832–3839. 10.1016/j.molimm.2008.05.022.

61. Waterhouse, A., Bertoni, M., Bienert, S., Studer, G., Tauriello, G., Gumienny, R., Heer, F.T., de Beer, T.A.P., Rempfer, C., Bordoli, L., et al. (2018). SWISS-MODEL: homology modelling of protein structures and complexes. Nucleic Acids Res 46, W296–w303. 10.1093/nar/gky427.

62. Kong, L., Lee, D.E., Kadam, R.U., Liu, T., Giang, E., Nieusma, T., Garces, F., Tzarum, N., Woods, V.L., Jr., Ward, A.B., et al. (2016). Structural flexibility at a major conserved antibody target on hepatitis C virus E2 antigen. Proc. Natl. Acad. Sci. U. S. A. 113, 12768–12773. 10.1073/pnas.1609780113.

